# Porcine ANTXR1, Heparan Sulfate and Neu5Gc act as entry factors for Seneca Valley virus invasion

**DOI:** 10.1101/2022.06.14.496051

**Authors:** Wenda Tang, Yanchao Wang, Xiaolan Qi, Fengxing Gu, Kangli Li, Haitang Han, Xuguang Du, Zixiang Zhu, Sen Wu, Yaofeng Zhao, Haixue Zheng

## Abstract

Seneca Valley virus (SVV) disease is a newly emerging infectious disease of pigs caused by SVV, which seriously endangers the pig industry. This study was set out to identify the essential host factors required for SVV entering porcine cells. Using a CRISPR/Cas9 library containing 93,859 sgRNAs that were designed to target approximately 22,707 porcine genes, we generated mutated porcine cell libraries, which were subjected to SVV challenge for enrichment of cells resistant to SVV infection. These resistant cells were subsequently analyzed to identify genes essential for SVV infection. We demonstrated that ANTXR1, a type I transmembrane protein encoded by *ANTXR1*, heparan sulfate (HS), glycosaminoglycans modified by acetylation and sulfation of HS2ST1, and Neu5Gc, a non-human sialic acid catalyzed by CMAH, were the essential host factors for SVV entry into porcine cells. These results will be helpful to elucidate the pathogenesis of SVV and the development of prevention and control measures.

## Introduction

Seneca Valley virus (SVV), belongs to the *Senecavirus* genus in the *picornaviridae* family, is responsible for a porcine idiopathic vesicular disease. Since 2014, SVV has been confirmed to be the causative agent of a newly emerging swine epidemic in the US ^1–4^, Brazil ^5–8^ and China ^9, 10^. To date, the global outbreak of SVV in swine has caused a sharp decline in the production of neonatal piglets and significant economic losses. However, no licensed vaccine or antiviral therapy is available yet, highlighting an urgent need for basic research on SVV.

Viruses rely on the host to complete their life cycles. Host cell entry factors play undoubtedly key roles determining the viral host range, tissue tropism, and viral pathogenesis. A thorough comprehension of interaction between host factors and virus will be the key to assess the impact of virus and will be helpful for finding better preventive and therapeutic tools. For example, as angiotensin-converting enzyme 2 (ACE2) was identified as the specific host receptor of severe acute respiratory syndrome coronavirus (SARS-CoV) ^11^, engineered soluble ACE2 has been used to compete with host receptor and thus prevent SARS-CoV-2 entry ^12^. In addition, gene editing of host factors has recently been shown to be an effective way defending against viruses in animal husbandry ^13^. For instance, CD163 serves as an uncoating receptor of porcine reproductive and respiratory syndrome virus (PRRSV). Based on understanding the key domains of CD163 interacting with PRRSV, pigs with precise gene editing of CD163 exhibited a full resistance to the porcine reproductive and respiratory syndrome (PRRS) ^14, 15, 16^.

Isolated in the cell culture of human fetal retinoblasts in 2002 ^17^, SVV was initially reported as a nonpathogenic oncolytic virus, as SVV showed high specificity for some human tumor cells rather than normal tissues. The anthrax toxin receptor-1 (ANTXR1) of human origin, alternatively named tumor endothelial marker 8 (TEM8) highly expressing in some cancer tissue as a tumor-specific endothelial marker, was previously identified as a SVV receptor in neuroendocrine cancer ^18^. It explained the tropism of SVV to human tumor cells. However, the essential host factors for SVV entry are yet to be determined in pigs.

CRISPR/Cas9-based loss-function libraries have recently been widely used for characterization of host factors associated with virus infections, such as SARS-CoV-2 ^19^, Rift Valley fever virus (RVFV) ^20^, and alphaviruses ^21^. In this work, a genome-wide CRISPR/Cas9 single-guide RNA library was designed for the pig and subsequently used to screen host factors for SVV susceptibility, and the functional role of the identified genes in mediating SVV infection was investigated to clarify the mechanism underlying infection, which allowed insights into prevention and treatment of the disease.

## Results

### A CRISPR/Cas9 mediated genome-wide knockout library was generated using IBRS-2 cells

To identify host factors required for SVV infection, we constructed a genome-scale loss-of-function library containing 93,859 CRISPR single-guide RNAs (sgRNAs), targeting 22,707 genes in the swine genome. To guarantee the targeting efficiency and to avoid the effects of off-targeting, we designed approximately five sgRNAs for each gene, and the target sites were mainly chosen in the first exon of the genes to improve the efficiency of the sgRNA. In addition, one non-targeting sgRNAs was included as a negative control. We subsequently integrated an all-in-one expression vector to simultaneously deliver Cas9, a sgRNA, and a puromycin resistance gene into the target cells (**Fig. 1a**). We measured that the 92,918 designed sgRNA oligonucleotides were successfully cloned into the plasmid library, and the relative abundance difference between the highest 10% sgRNA and the lowest 10% sgRNA was within 15 fold (**Fig. 1b**). The plasmid pools were packaged into lentivirus in HEK293-T cells. 1.2×10^8^ IBRS-2 cells, which are permissive to SVV, were transduced with the lentivirus library at 0.2 MOI to ensure that the majority of cells harbored one sgRNA (**Fig. 1c**). The cells were cultured for 7 days after a puromycin selection was completed to make sure that the protein functions were destroyed. The genomic DNA was isolated and the sgRNA regions were amplified by PCR. Deep sequencing results showed that 85,151 guide RNAs were retained in genome, covering about 91% of the original sequences (**Fig. 1d**).

**Fig 1.**
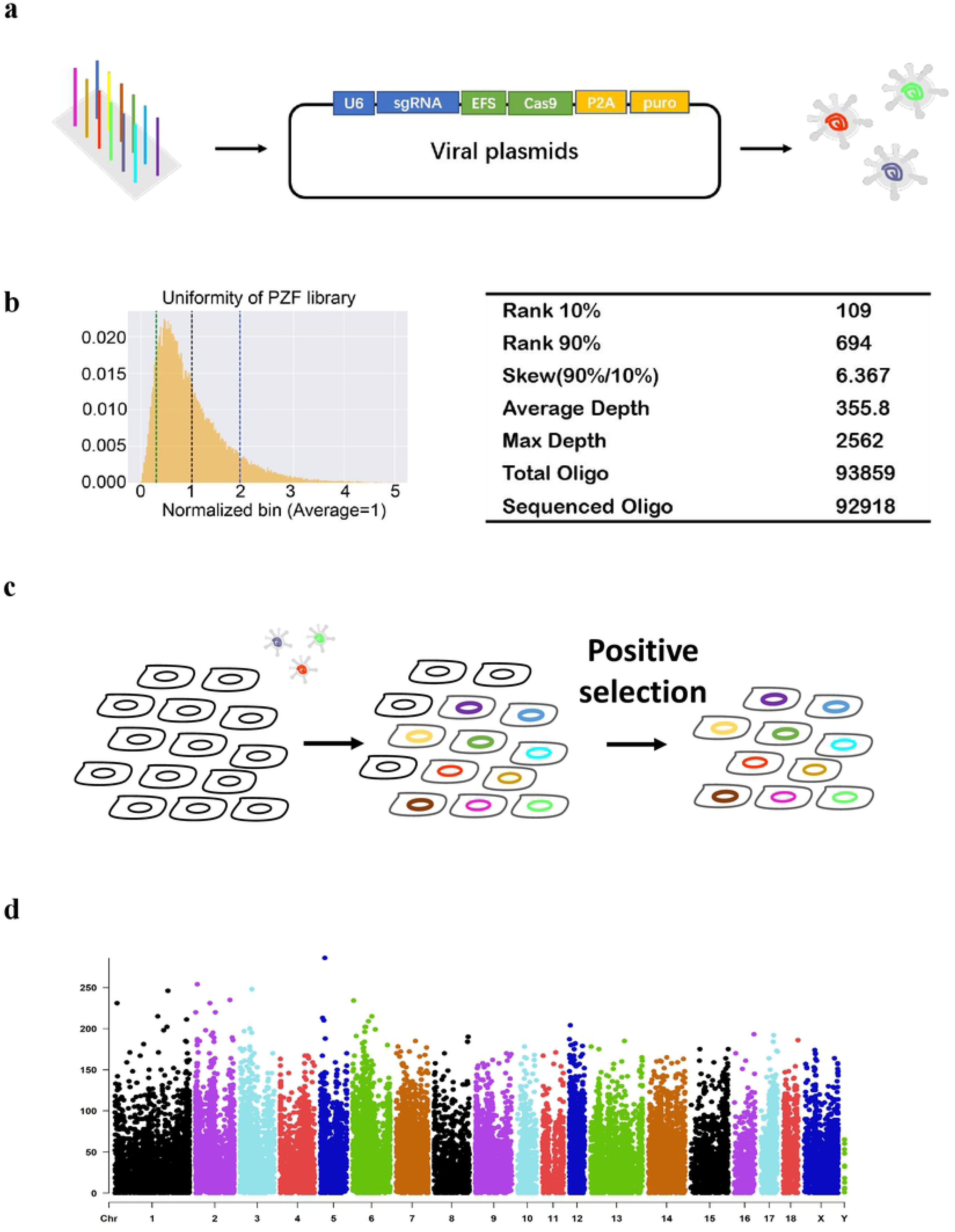
Construction of porcine genome-wide mutant cell library. **a Schematic diagram of plasmids library construction.** Oligo array synthesized sgRNAs were cloned into the plasmids, then packaged to lentivirus in HEK293-T cells. **b Quality detection of plasmid library.** The abscissa represents each oligo and the ordinate represents read counts after normalization. The specific sequencing values are presented in the table on the right. **c Schematic diagram of cell mutation library construction.** Lentiviral pools were transduced into cells, positive cell clones were selected by puromycin. **d Distribution of mutant genes on genome.** The abscissa represents each chromosome and the ordinate represents read counts.

### A candidate gene list for Seneca Valley virus infection was obtained by CRISPR/Cas9 screen

The selected cell pools were challenged with SVV at 1.0 MOI for 24 h, with a clear cytopathic effect (CPE). The DNA was extracted from the surviving cells for PCR amplification and then subjected to next-generation sequencing (NGS) and analysis. A significant reduction in the diversity of sgRNAs in the surviving cells was observed, reflecting that the sgRNAs targeted essential genes (**Fig. 2a, Supplementary Fig. 1a, b**).

**Fig 2.**
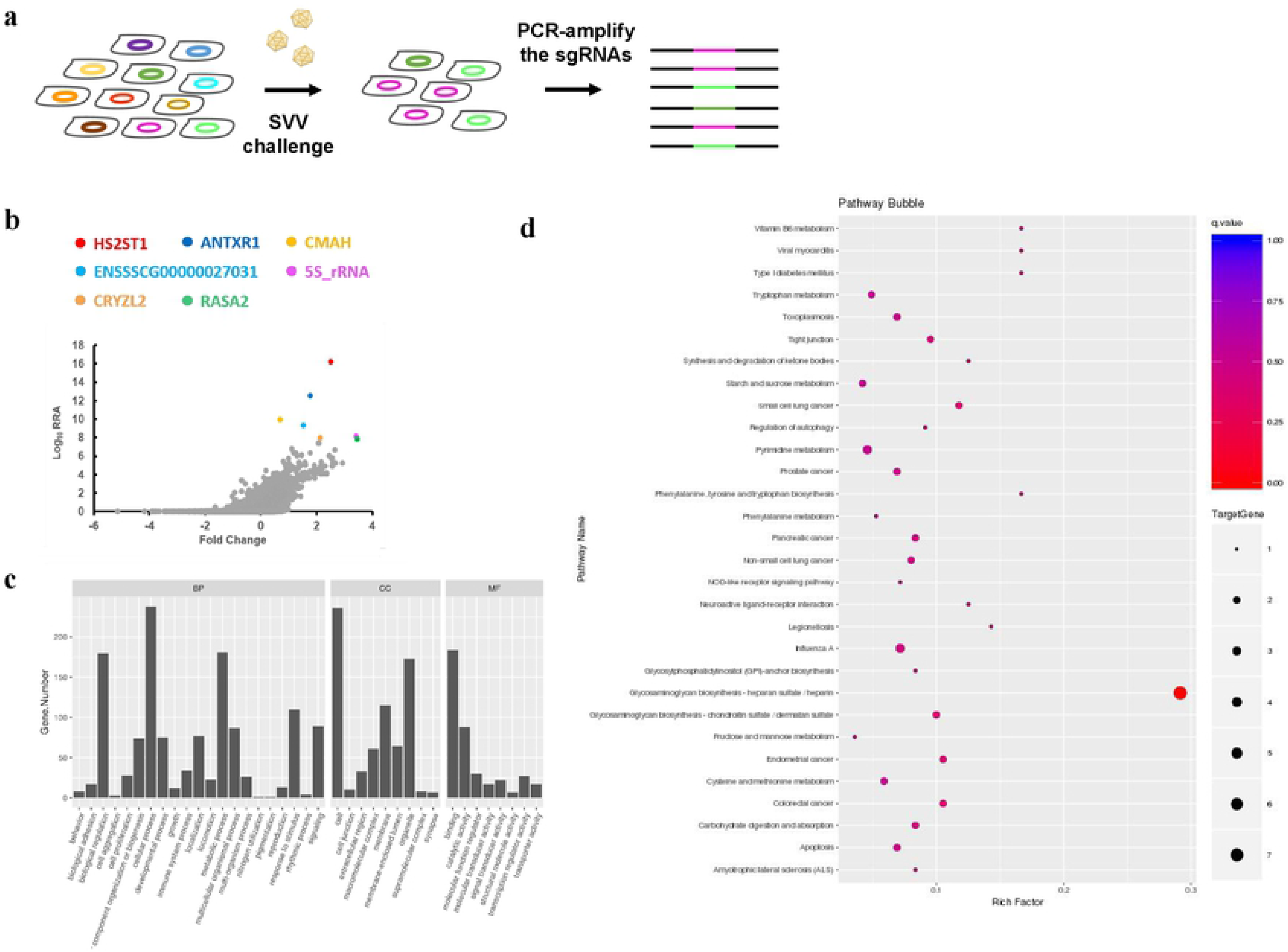
SVV resistance related genes were obtained by screening. **a Schematic diagram of SVV-resistant cell library construction.** Cell mutation library were challenged with SVV (MOI = 1) for 24 h. Genomic DNA was extracted from the surviving cells. The sgRNA sequences were amplified and subjected to next generation sequencing. **b Ranking of candidate genes.** The top seven hits are shown as different colors. The abscissa represents log fold change and the ordinate represents log Robust Rank Aggreg (RRA). **c Gene Ontology (GO) analysis for the enriched gene targets.** The histogram shows the names of the top 30 GO terms and the corresponding number of genes. According to the GO classification, the Classes are divided into: BP (biological process,), MF (molecular function) and CC (cellular component). **d Kyoto Encyclopedia of Genes and Genomes (KEGG) analysis for the enriched gene targets.** The scatter plot is a graphical representation of the KEGG enrichment analysis. KEGG enrichment was measured by Rich Factor, q-value, and the number of genes enriched on this pathway. In this study, the 30 pathways with the most significant enrichment were selected and displayed in this figure.

The top 7 genes that showed the highest screen scores in three independent experiments are shown in **Fig. 2b** and **Table S1**: ENSSSCG00000022032 (heparan sulfate 2-O-sulfotransferase 1, *HS2ST1*), ENSSSCG00000008340 (ANTXR cell adhesion molecule 1, *ANTXR1*), ENSSSCG00000001099 (cytidine monophospho-N-acetylneuraminic acid hydroxylase, *CMAH*), ENSSSCG00000027031, ENSSSCG00000018405, ENSSSCG00000028377 (crystallin zeta like 1, *CRYZL1*), and ENSSSCG00000011672 (RAS p21 protein activator 2, *RASA2*). We further performed Gene Ontology (GO) analysis (**Fig. 2c**) and Kyoto Encyclopedia of Genes and Genomes (KEGG) analysis (**Fig. 2d**) on the data. From the KEGG analysis, we found that the glycosaminoglycan synthesis pathway not only showed the most significant enrichment but also contained the most abundant pathway. Of note, the genes from the enriched list highly likely to be involved in viral entry were: *ANTXR1*, encoding a type I transmembrane protein; *HS2ST1*, relating to heparan sulfate (HS) modification and *CMAH*, responsible for catalytic synthesis of sialic acid.

### Knockout of *ANTXR1* leads to a significant reduction of Seneca Valley virus **permissivity**

The second most significantly enriched gene in the SVV screen library were found to be the *ANTXR1* gene, encoding a single-pass cell surface protein consisting of an intracellular region, transmembrane region, and an extracellular region with a von Willebrand Factor A (vWA) domain ^22, 23, 24^. To validate the function of ANTXR1 in SVV infection in pigs, knockout cells (PK-15 and CRL-2843) were generated using the CRISPR/Cas9 system (**Supplementary Fig. 2a, b**). The *ANTXR1* KO cell lines were infected with SVV for 12 h. As expected, viral replication was significantly decreased as shown by qPCR (**Fig. 3a, e**). A reduction in the protein level was also confirmed by Western blotting (**Fig. 3b, f**). Cytopathic effects (CPE) were not observed in CRL-2843 after *ANTXR1* knockout (**Supplementary Fig. 3a**), and no virus was observed in the knockout (KO) cell lines using immunofluorescence staining (**Fig. 3g, Supplementary Fig. 3b**), indicating that the cell lines had gained protection against the SVV infection. Moreover, exogenous expression of the ANTXR1 restored the susceptibility of gene knockout lines to viral permissivity, suggesting that the antiviral phenotype of cells is indeed caused by the deletion of *ANTXR1* (**Fig. 3c, d**).

**Fig 3.**
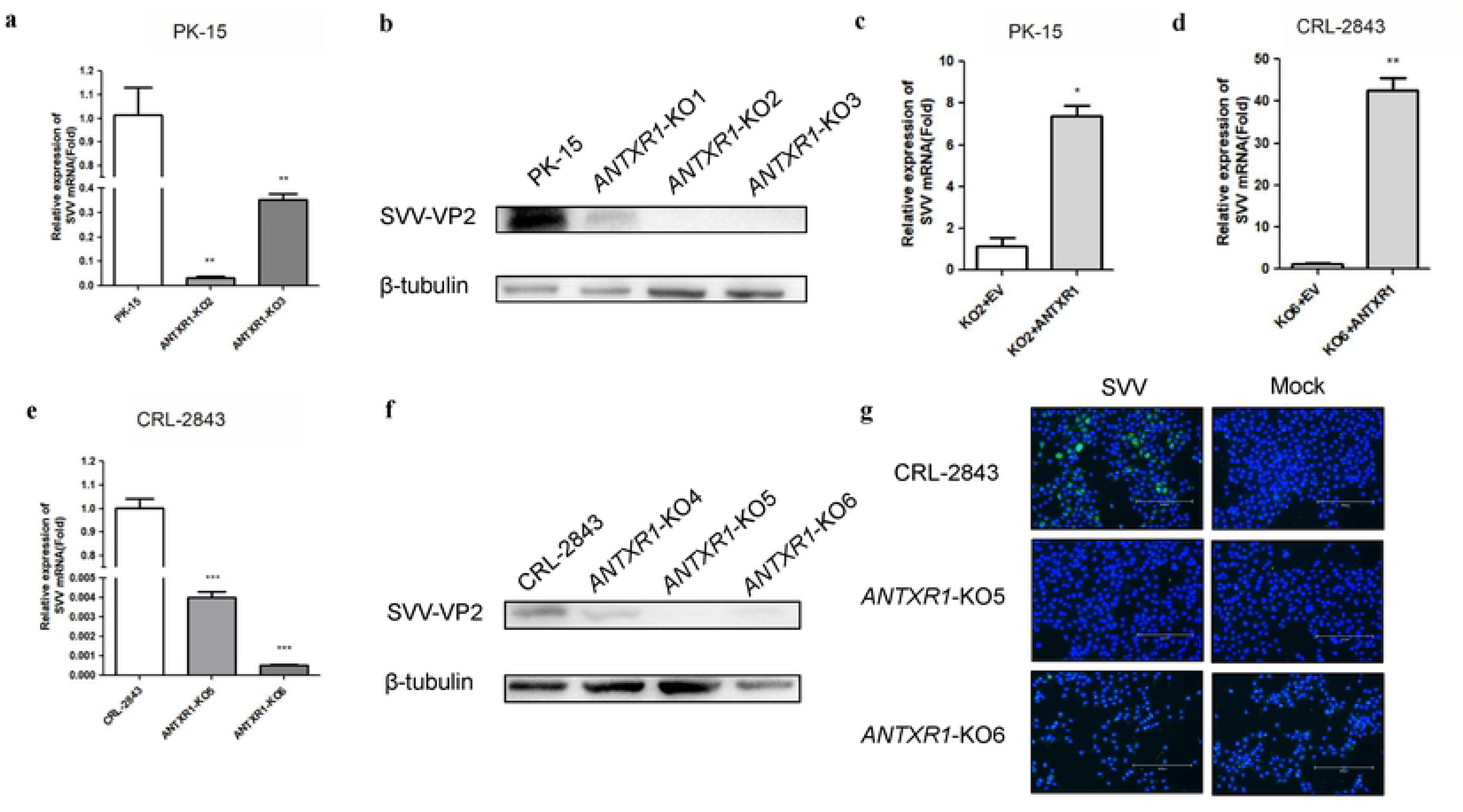
Knockout of *ANTXR1* significantly reduced SVV infection. **a Quantitative analysis of SVV RNA in *ANTXR1*-knockout and wild type PK-15 cells.** Wild type PK-15 cells and *ANTXR1* KO cells were cultured in 12 well plates, and infected with SVV (MOI = 1). At 12 hpi (hours post infection), SVV mRNAs was determined with qPCR assay. **b Western blotting detection of SVV-VP2 in *ANTXR1-*knockout and wild PK-15 type cells.** Wild type PK-15 cells and *ANTXR1* KO cells were cultured in six well plates, and infected with SVV (MOI = 1). At 12 hpi, the cells were collected for Western blotting. **c d Quantitative analysis of SVV RNA in *ANTXR1-*knockout cells after transfection.** The *ANTXR1* KO cells were transfected with 3 μg Myc–ANTXR1-expressing plasmid. At 48 hpt (hours post transfection), SVV was added (MOI = 1) for 12 h. The same amount of empty vector was used in the transfection process. SVV mRNAs were determined with qPCR assay. **e Quantitative analysis of SVV RNA in *ANTXR1*-knockout and wild type CRL-2843 cells.** Wild type CRL-2843 cells and *ANTXR1* KO cells were cultured in 12 well plates, and infected with SVV (MOI = 1). At 12 hpi, SVV mRNAs was determined with qPCR assay. **f Western blotting detection of SVV-VP2 in *ANTXR1-*knockout and wild type CRL-2843 cells.** Wild type CRL-2843 cells and *ANTXR1* KO cells were cultured in six well plates, and infected with SVV (MOI = 1). At 12 hpi, the cells were collected for Western blotting. **g Indirect immunofluorescence assay of SVV-VP2 in *ANTXR1-*knockout and wild type CRL-2843 cells.** Wild type CRL-2843 cells and *ANTXR1* KO cells were cultured in 24 well plates, and infected with SVV (MOI = 1). At 8 hpi, VP2 protein expression (green) was detected by indirect immunofluorescence assay. Cell nuclei were stained with a NucBlue Live ReadyProbe (blue). Scale bars: 300 μm. Data are means ± SD of triplicate samples. \**P* < 0.05, \*\**P* < 0.01, \*\*\**P* < 0.001 (two-tailed Student’s t-test)

### ANTXR1 is the cellular receptor for Seneca Valley virus

To further analyze the potential role of pig ANTXR1 in SVV entry, we examined whether viral entry into the cells was inhibited in the absence of ANTXR1. Incubating with SVV at 4 °C for 1 h or 37 °C for 30 min respectively, *ANTXR1* KO cell lines displayed a significantly decreased viral attachment and internalization (**Fig. 4a, b**). Furthermore, ectopic expression of pig ANTXR1 enabled SVV attachment (**Fig. 4c**) and an 11-fold increase in virus infection for 24 h (**Fig. 4d**) in MDCK, a cell line which normally is not permissive for SVV infection.

**Fig 4.**
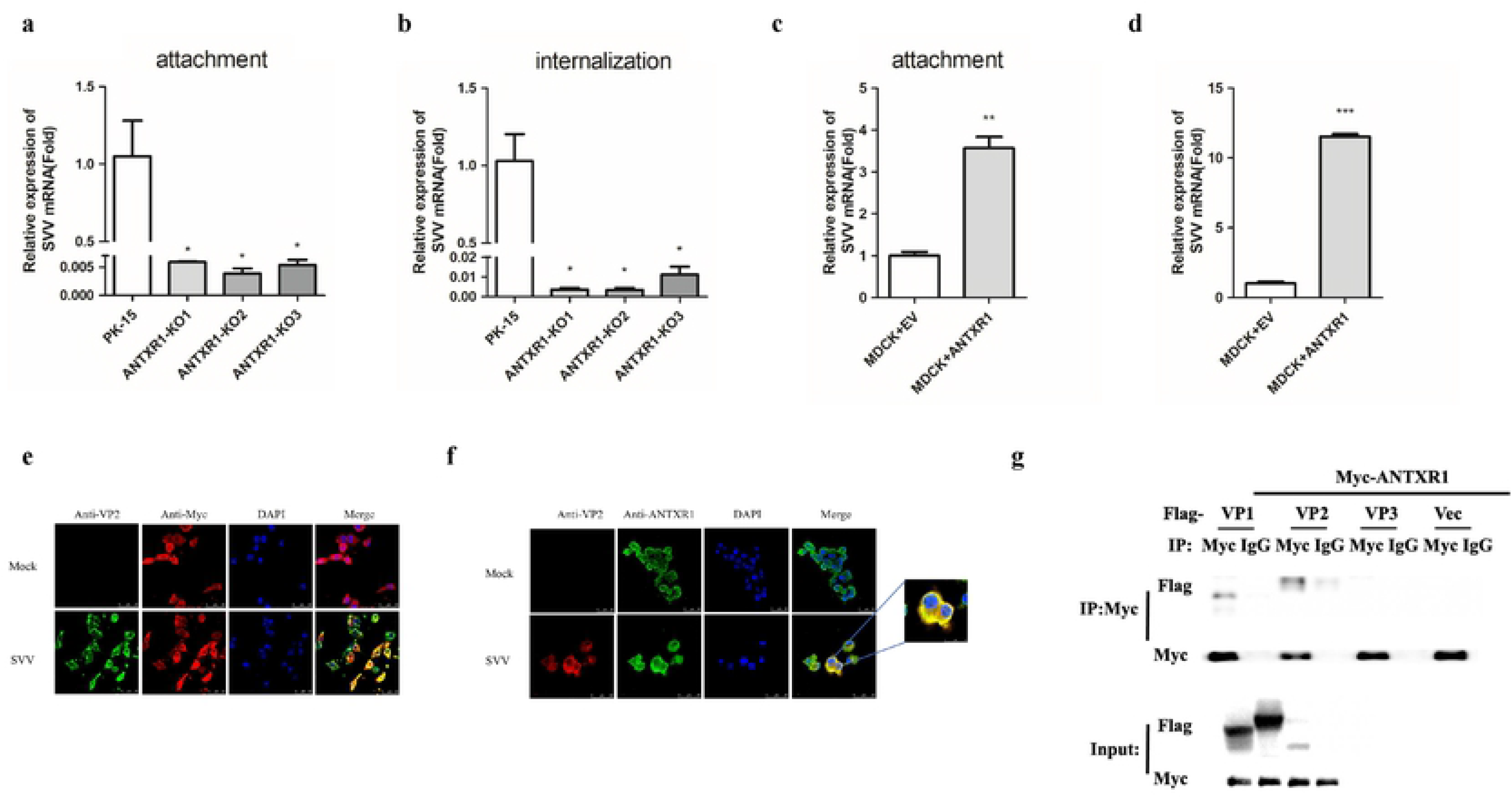
ANTXR1 affects the SVV enter cells by interacting with VP1 and VP2. **a Quantitative analysis of SVV RNA in *ANTXR1*-knockout and wild type PK-15 cells during the process of SVV attachment.** Wild type PK-15 cells and *ANTXR1* KO cells were cultured in 12 well plates, and infected with SVV (MOI = 10) at 4 °C for 1 h. SVV mRNAs was determined with qPCR assay. **b Quantitative analysis of SVV RNA in *ANTXR1*-knockout and wild type PK-15 cells during the process of SVV internalization.**Wild type PK-15 cells and *ANTXR1* KO cells were cultured in 12 well plates, and infected with SVV (MOI = 10) at 37 °C for 30 min. SVV mRNAs was determined with qPCR assay. **c Quantitative analysis of SVV RNA in MDCK cells during the process of SVV attachment.** MDCK cells were transfected with 3 μg Myc–ANTXR1-expressing plasmid. At 48 hpt, SVV was added (MOI = 10) at 4 °C for 1 h. SVV mRNAs was determined with qPCR assay. **d Quantitative analysis of SVV RNA in MDCK cells.** MDCK cells were transfected with 3 μg Myc–ANTXR1-expressing plasmid. At 48 hpt, SVV was added (MOI = 10) for 24 h. SVV mRNAs was determined with qPCR assay. **e Observation of the co-localization of porcine ANTXR1 and SVV in HEK-293T cells by laser confocal experiment.** HEK293-T cells were transfected with 3 μg Myc–ANTXR1-expressing plasmid, then SVV was added 24 h later (MOI = 1) for 9 h. Images were obtained by confocal microscopy using a 100× objective. Scale bars: 50 μm. **f Observation of the co-localization of ANTXR1 and SVV in IBRS-2 cells by laser confocal experiment.** IBRS-2 cells were infected with SVV (MOI = 1) for 9 h. Images were obtained by confocal microscopy using a 100× objective. Scale bars: 50 μm and 25 μm. **g Coimmunoprecipitation.** HEK293-T cells were cultured in 10-cm dishes and transfected with 8 μg Myc–ANTXR1, 8 μg viral structural protein with a Flag tag expressing plasmid. The cells were collected for co-immunoprecipitated 24 h later. Data are means ± SD of triplicate samples. \**P* < 0.05, \*\**P* < 0.01, \*\*\**P* < 0.001 (two-tailed Student’s t-test)

The pcDNA3.1 vector containing pig ANTXR1 was transfected into HEK293-T cells and SVV was added 24 h later. SVV-ANTXR1 co-localization was observed after 8 h (**Fig. 4e**). We also tested the co-localization of endogenous ANTXR1 and SVV in IBRS-2 cells (**Fig. 4f**). To investigate interactions between ANTXR1 and SVV structural proteins, HEK293-T cells were transfected using a Myc–ANTXR1 expressing vector, with Flag-tagged viral structural proteins VP1, VP2, VP3 expressing plasmids, respectively. Analyzed by Western blotting, we found that ANTXR1 interacts with VP1 and VP2 (**Fig. 4g**). These results supported the hypothesis that pig ANTXR1 is the cellular receptor for SVV infection.

### Heparan sulfate is associated with Seneca Valley virus infection

A large number of genes encoding enzymes involved in heparan sulfate (HS) synthesis and modification were enriched (*p*<0.01) in the candidate gene list, such as *B3GAT3* (ranking number 2211), *EXT1* (ranking number 37), *EXT2* (ranking number 2492), *NDST2* (ranking number 3559), *NDST3* (ranking number 257), *HS2ST1* (ranked at the very top) and *HS3ST5* (ranking number 1295). Most strikingly, we found that the HS synthesis pathway was upregulated in the KEGG analysis in the transcriptome sequences of wild-type cells after SVV infection, including *EXTL1*, *EXT1*, *HS3ST1* and *HS3ST3B1* (**Fig. 5a, Supplementary Fig. 4**).

**Fig 5.**
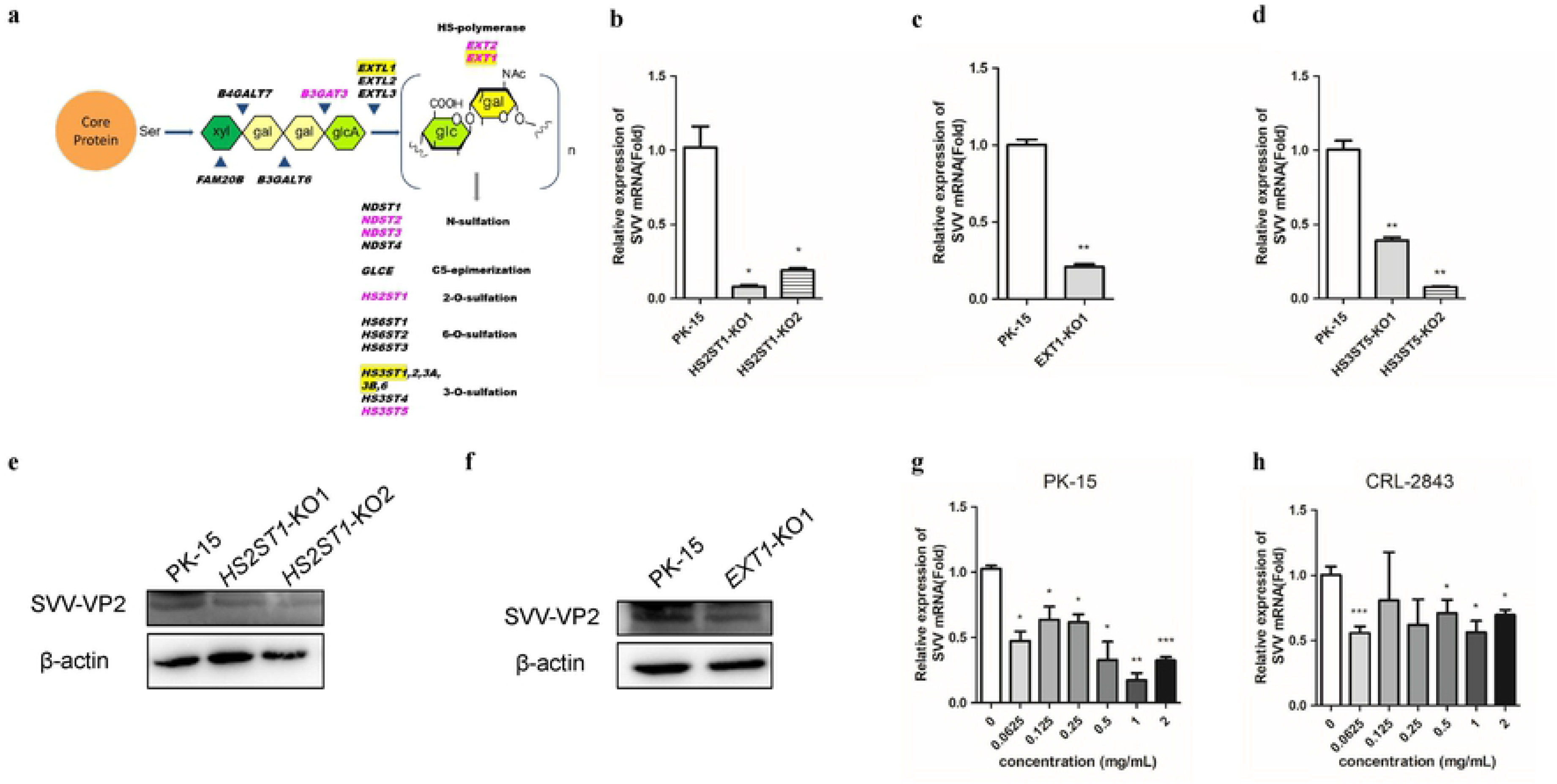
HS is the essential for SVV republication. **a Heparan sulfate (HS)/heparin biosynthetic pathways.** Enriched genes were significantly enriched in this screen are indicated in pink. Significantly upregulated genes after SVV infection in transcriptome analysis of wild-type PK-15 cells are indicated in yellow background. **b Quantitative analysis of SVV RNA in *HS2ST1*-knockout and wild type cells.** Wild type PK-15 cells and *HS2ST1* KO cells were cultured in 12 well plates, and infected with SVV (MOI = 1). At 12 hpi, SVV mRNAs was determined with qPCR assay. **c Quantitative analysis of SVV RNA in *EXT1*-knockout and wild type cells.** Wild type PK-15 cells and *EXT1* KO cells were cultured in 12 well plates, and infected with SVV (MOI = 1). At 12 hpi, SVV mRNAs was determined with qPCR assay. **d Quantitative analysis of SVV RNA in *HS3ST5*-knockout and wild type cells.** Wild type PK-15 cells and *HS3ST5* KO cells were cultured in 12 well plates, and infected with SVV (MOI = 1). At 12 hpi, SVV mRNAs was determined with qPCR assay. **e Western blotting detection of SVV-VP2 in *HS2ST1*-knockout and wild type cells.** Wild type PK-15 cells and *HS2ST1* KO cells were cultured in six well plates, and infected with SVV (MOI = 1). At 12 hpi, the cells were collected for Western blotting. **f Western blotting detection of SVV-VP2 in *EXT1*-knockout and wild type cells.** Wild type PK-15 cells and *EXT1* KO cells were cultured in six well plates, and infected with SVV (MOI = 1). At 12 hpi, the cells were collected for Western blotting. **g Quantitative analysis of SVV RNA in PK-15 cells with different concentrations of heparin sodium treatment.** PK-15 cells were incubated with soluble heparin sodium for 30 min, then infected with SVV for 12 h. SVV mRNAs was determined with qPCR assay. **h Quantitative analysis of SVV RNA in CRL-2843 cells with different concentrations of heparin sodium treatment.** CRL-2843 cells were incubated with soluble heparin sodium for 30 min, then infected with SVV for 12 h. SVV mRNAs was determined with qPCR assay. Data are means ± SD of triplicate samples. \**P* < 0.05, \*\**P* < 0.01, \*\*\**P* < 0.001 (two-tailed Student’s t-test)

Therefore, to investigate whether HS serves as an essential factor for SVV infection, we performed mutations in *EXT1*, *HS2ST1* and *HS3ST5* (**Supplementary Fig. 5a-c**) and then measured the effect on SVV infection. The relative amount of SVV mRNA was around 5-fold lower in *EXT1* KO cells compared to wild type PK-15 cells (**Fig. 5b**), while the relative mRNA of SVV was reduced 5-fold to 10-fold in the *HS2ST1* KO cells (**Fig. 5c**), 2-fold to 12-fold in the *HS3ST5* KO cells compared to wild type PK-15 cells (**Fig. 5d**), reflecting that the KO cell lines gained resistance to SVV infection. Western blotting also confirmed the low level of SVV-VP2 protein in KO cells as compared to wild type cells (**Fig. 5e, f**).

Soluble heparin sodium, an analogue of heparan sulfate, is usually used as a HS competitive binding reagent. SVV was added to PK-15 cells after treatment with different concentrations of soluble heparin sodium for 1 h, which reduced viral infection substantially at 0.5 mg/mL and higher concentrations (**Fig. 5g**). Under the same condition, SVV infection was also reduced by soluble heparin sodium in CRL-2843 cells (**Fig. 5h**).

### Heparan sulfate affects virus attachment

To determine whether HS could affect SVV binding to the cell surface, the ability of SVV attachment and internalization were further analyzed in *HS2ST1* and *EXT1*KO cell lines. The SVV attachment and internalization were significantly reduced in the *HS2ST1* KO cells compared with the wild type PK-15 cells (**Fig. 6a, b**). The same results were also observed in the *EXT1* KO cells (**Fig. 6c, d**). In addition, heparin sodium also very potently reduced the SVV attachment and internalization (**Fig. 6e-h**). These data further demonstrated that HS serves as an adhesion factor for SVV.

**Fig 6.**
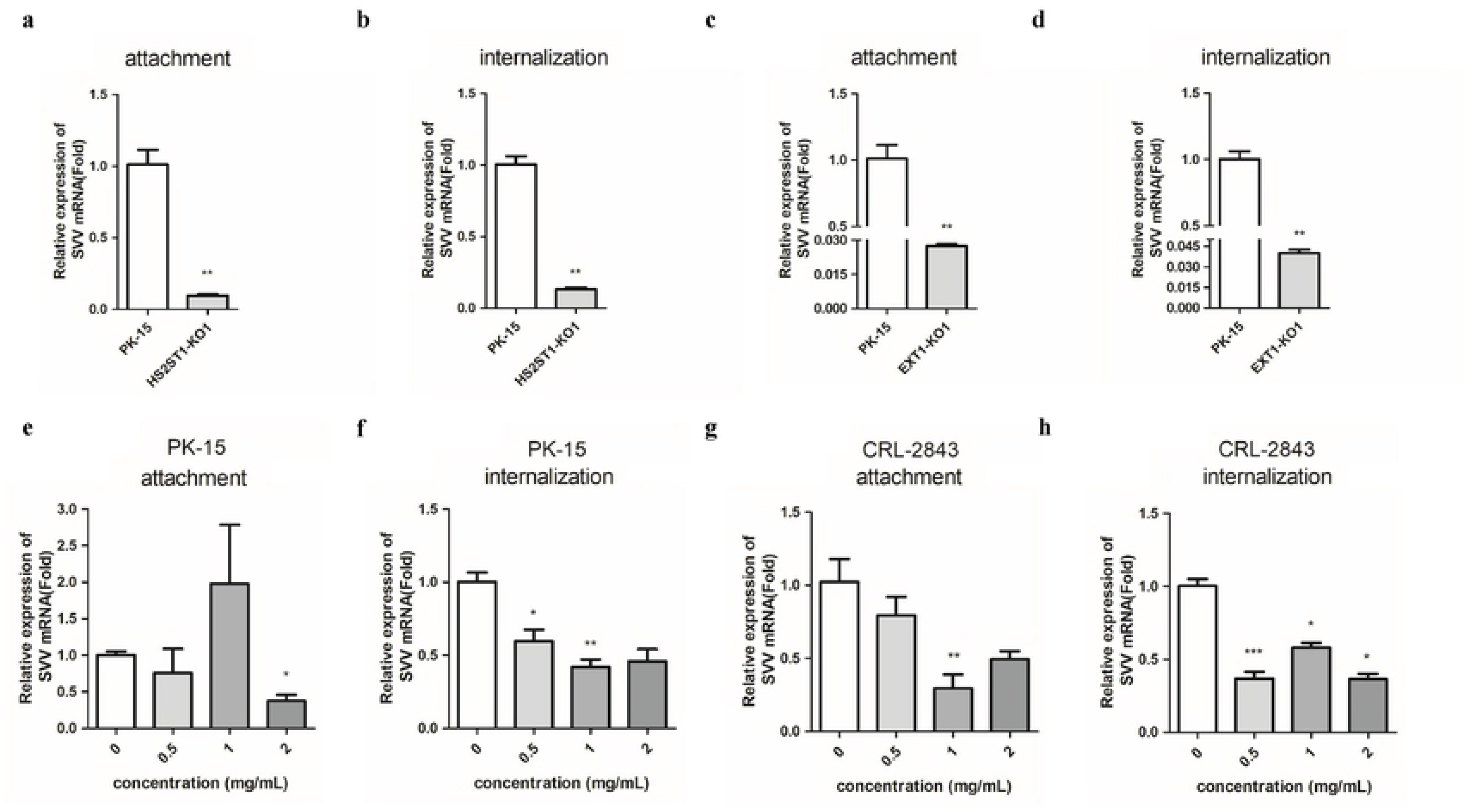
HS help to SVV attachment. **a Quantitative analysis of SVV RNA in *HS2ST1*-knockout and wild type PK-15 cells during the process of SVV attachment.** Wild type PK-15 cells and *HS2ST1* KO cells were cultured in 12 well plates, and infected with SVV (MOI = 10) at 4 °C for 1 h. SVV mRNAs was determined with qPCR assay. **b Quantitative analysis of SVV RNA in *HS2ST1*-knockout and wild type PK-15 cells during the process of SVV internalization.** Wild type PK-15 cells and *HS2ST1* KO cells were cultured in 12 well plates, and infected with SVV (MOI = 10) at 37 °C for 30 min. SVV mRNAs was determined with qPCR assay. **c Quantitative analysis of SVV RNA in *EXT1*-knockout and wild type PK-15 cells during the process of SVV attachment.** Wild type PK-15 cells and *EXT1* KO cells were cultured in 12 well plates, and infected with SVV (MOI = 10) at 4 °C for 1 h. SVV mRNAs was determined with qPCR assay. **d Quantitative analysis of SVV RNA in *EXT1*-knockout and wild type PK-15 cells during the process of SVV internalization.** Wild type PK-15 cells and *EXT1* KO cells were cultured in 12 well plates, and infected with SVV (MOI = 10) at 37 °C for 30 min. SVV mRNAs was determined with qPCR assay. **e Quantitative analysis of SVV RNA in PK-15 cells with different concentrations of heparin sodium treatment during the process of SVV attachment.** PK-15 cells were incubated with soluble heparin sodium for 30 min, then infected with SVV at 4 °C for 1 h. SVV mRNAs was determined with qPCR assay. **f Quantitative analysis of SVV RNA in PK-15 cells with different concentrations of heparin sodium treatment during the process of SVV internalization.** PK-15 cells were incubated with soluble heparin sodium for 30 min, then infected with SVV at 37 °C for 30 min. SVV mRNAs was determined with qPCR assay. **g Quantitative analysis of SVV RNA in CRL-2843 cells with different concentrations of heparin sodium treatment during the process of SVV attachment.** CRL-2843 cells were incubated with soluble heparin sodium for 30 min, then infected with SVV at 4 °C for 1h. SVV mRNAs was determined with qPCR assay. **h Quantitative analysis of SVV RNA in CRL-2843 cells with different concentrations of heparin sodium treatment during the process of SVV internalization.** CRL-2843 cells were incubated with soluble heparin sodium for 30 min, then infected with SVV at 37 °C for 30 min. SVV mRNAs was determined with qPCR assay. Data are means ± SD of triplicate samples. \**P* < 0.05, \*\**P* < 0.01, \*\*\**P* < 0.001 (two-tailed Student’s t-test)

### Functional validation of CMAH in SVV infection

To validate the function of CMAH in SVV infection, knockout cells (IBRS-2) were generated using the CRISPR/Cas9 system (**Supplementary Fig. 6**). The *CMAH* KO cell lines were infected with SVV for 12 h, and the relative amount of mRNA encoding SVV was about 4-fold lower in *CMAH* KO cells compared to wild type IBRS-2 cells (**Fig. 7a**). Western blotting results indicated that viral VP2 was decreased significantly (**Fig. 7b**). Moreover, another *CMAH* KO cell line (IBRS-2) also had a similar phenotype (**Fig. 7c**). As demonstrated in **Fig. 7d**, ectopic expression of pig CMAH restored the susceptibility of *CMAH* KO lines, suggesting that SVV infection depends on CMAH.

**Fig 7.**
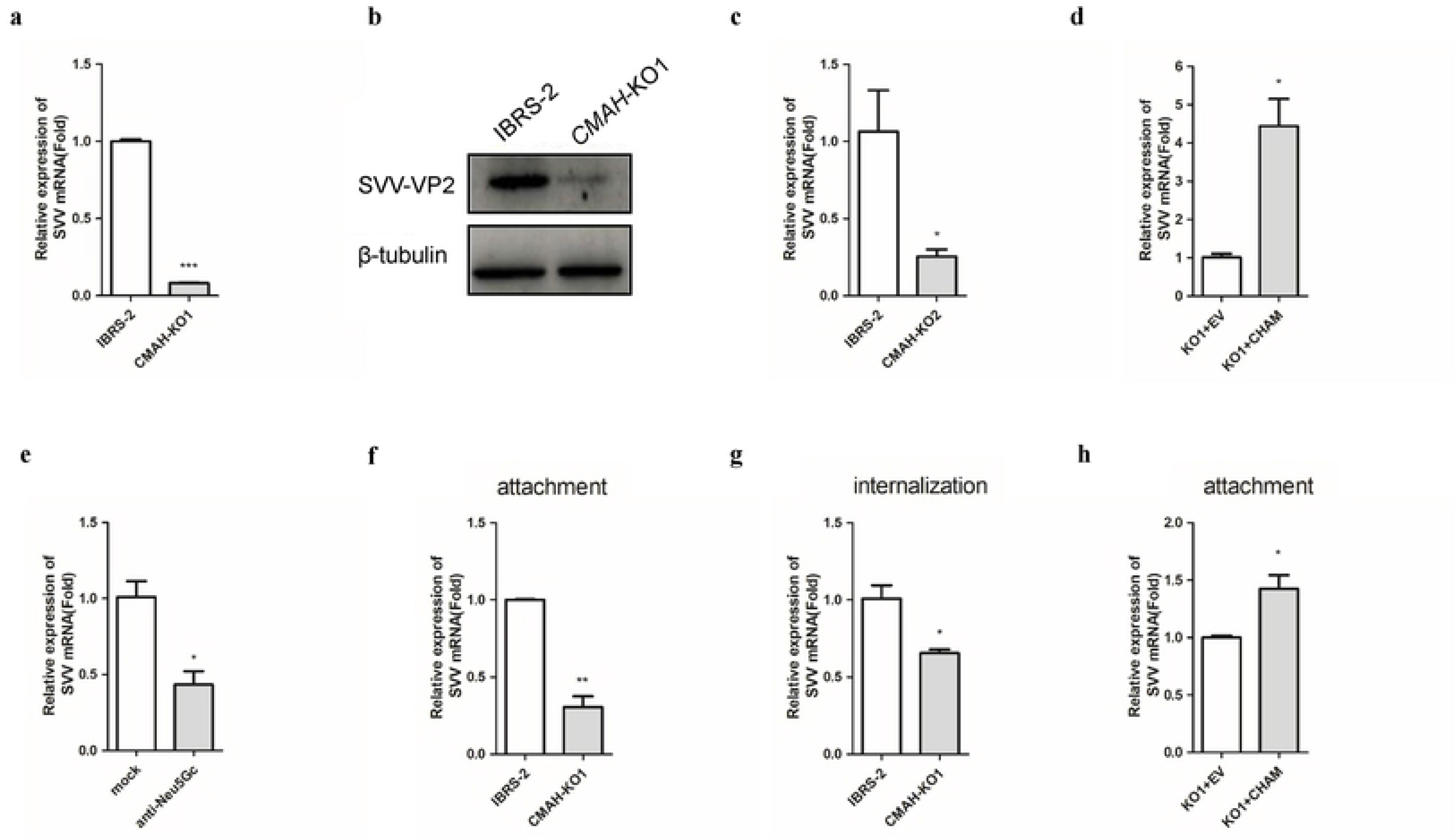
**S**eneca Valley virus entry depends on CMAH. **a Quantitative analysis of SVV RNA in *CMAH*-KO1 and wild type cells.** Wild type IBRS-2 cells and *CMAH* KO1 cells were cultured in 12 well plates, and infected with SVV (MOI = 1). At 12 hpt, SVV mRNAs was determined with qPCR assay. **b Western blotting detection of SVV-VP2 in *CMAH*-KO1 and wild type cells.** Wild type IBRS-2 cells and *CMAH* KO cells were cultured in six well plates, and infected with SVV (MOI = 1). At 12 hpt, the cells were collected for western blotting. **c Quantitative analysis of SVV RNA in *CMAH-*KO2 and wild type cells.** Wild type IBRS-2 cells and *CMAH* KO2 cells were cultured in 12 well plates, and infected with SVV (MOI = 1). At 12 hpt, SVV mRNAs was determined with qPCR assay. **d Quantitative analysis of SVV RNA in *CMAH-*knockout cells after transfection.** The *CMAH* KO cells were transfected with 2 μg Myc–CMAH-expressing plasmid. At 24 hpt, SVV was added (MOI = 1) for 24 h. The same amount of empty vector was used in the transfection process. SVV mRNAs were determined with qPCR assay. **e Anti-Neu5Gc Antibody blocking assay.** Anti-Neu5Gc antibody were preincubated at 1:200 dilution with IBRS-2 cells for 1h, then SVV was added (MOI = 1). After 1h, media was replaced by DMEM. At 12 hpi, SVV mRNAs was determined with qPCR assay. **f Quantitative analysis of SVV RNA in *CMAH*-knockout and wild type IBRS-2 cells during the process of SVV attachment.** Wild type IBRS-2 cells and *CMAH* KO cells were cultured in 12 well plates, and infected with SVV (MOI = 10) at 4 °C for 1h. SVV mRNAs was determined with qPCR assay. **g Quantitative analysis of SVV RNA in *CMAH*-knockout and wild type IBRS-2 cells during the process of SVV internalization.** Wild type IBRS-2 cells and *CMAH* KO cells were cultured in 12 well plates, and infected with SVV (MOI = 10) at 37 °C for 30 min. SVV mRNAs was determined with qPCR assay. **h Quantitative analysis of SVV RNA in *CMAH-*knockout cells after transfection during the process of SVV attachment.** The *CMAH* KO cells were transfected with 2 μg Myc–CMAH-expressing plasmid. At 48 hpt, SVV was added (MOI = 10) at 4 °C for 1h. SVV mRNAs were determined with qPCR assay. Data are means ± SD of triplicate samples. \**P* < 0.05, \*\**P* < 0.01, \*\*\**P* < 0.001 (two-tailed Student’s t-test)

### Seneca Valley virus entry depends on CMAH

CMAH is the key enzyme to synthesize Neu5Ac into Neu5Gc. Previous studies have shown that Neu5Gc is a cell surface receptor for influenza virus. We thus measured viral attachment, viral entry, and internalization to determine how CMAH affects SVV infection. *CMAH* KO cells showed significantly reduced SVV attachment and internalization (**Fig. 7e, f**). The ectopic expression of pig CMAH increased SVV attachment (**Fig. 7g**), whereas the antibody of Neu5Gc reduced SVV infection significantly (**Fig. 7h**). These results demonstrated that SVV entry depends on Neu5Gc.

## Discussion

We applied CRISPR/Cas9 for genome-wide screening in porcine cells in order to elucidate the host factors determining the susceptibility to SVV infection, showing that ANTXR1, heparan sulfate and Neu5Gc play important roles in SVV entry.

ANTXR1 has been confirmed as a SVV receptor in selected human tumor cells. According to previous structural studies, the R88 site and D156 site of human ANTXR1 are critical amino acid residues interacting with SVV ^25, 26^. However, the corresponding sites in pig ANTXR1 mutate to other amino acids. Despite the difference, we demonstrated in this study that ANTXR1 also acts a receptor mediating SVV entry into porcine cells, and it physically interacts with VP1 and VP2 of SVV. As SVV infects pigs but not humans ^6^, this finding suggests that more complicated factors but not a single receptor determine the species tropism of SVV.

The conservation of *ANTXR1* among different species suggests that it may have other unknown important physiological functions. As reported, ANTXR1 may be associated with collagens binding and promotion of ECs migration and misaligned incisors are observed in adult *ANTXR1* KO mice ^27^. Mutations in *ANTXR1* cause progressive extracellular-matrix accumulation in patients with the GAPO syndrome, a complex phenotype consisting of growth retardation, alopecia, pseudoanodontia and progressive optic atrophy ^28^. Recently, *ANTXR1* knockout pigs were produced, and as expected, exhibited resistance to SVV infection, while these pigs also developed GAPO-like symptoms ^29^. Based on the above results, more explicit structural investigations on the SVV-ANTXR1 complex are warranted in order to design accurate editing sites that destroy the function of virus receptor while maintaining its normal physiological function ^30^.

Heparan sulfate (HS) is a highly sulfated polysaccharide, which is widely distributed on the surface of cell membranes, basement membranes and the extracellular matrix. Its synthesis is catalyzed by a series of synthases and modifying enzymes ^31^. Various structures and modifications lead to the complexity and diversity of biological properties of heparan sulfate, including cell adhesion, regulation of cell growth and proliferation, and development processes ^32, 33, 34^. Because of its anionic characteristics and high-density negative charge, heparan sulfate possesses the ability to interact with viruses. Heparan sulfate has been identified as receptor or co-receptor for certain types of viruses, such as SARS-CoV2 ^35^, lymphocytic choriomeningitis virus (LCMV) ^36^, and chikungunya virus ^37^. A series of HS synthetases and sulfation modifying enzymes were shown to be enriched in our experiment, and we demonstrated that heparan sulfate is also a receptor for SVV. By exogenous addition of soluble heparin sodium, competitive binding of heparan sulfate and heparin sodium to SVV was studied. We found that SVV invasion and infection were both impaired. This suggests that low-cost small molecule therapeutic drugs may block the combination of virus and HS, so as to reduce the virus infection in pigs.

Sialic acids constitute a 9-carbon monosaccharide family that is important for a wide variety of biological events. The predominant sialic acids in mammals are N-acetylneuraminic acid (Neu5Ac) and N-glycolylneuraminic acid (Neu5Gc), the latter is formed by hydroxylation by CMAH. The *CMAH*^−/−^ knockout mice model was previously used to investigate the effect of Neu5Gc on influenza virus infection, showing that Neu5Gc acted as a functional receptor ^38^. Porcine *CMAH* KO lessens the severity of the PEDV infection and delays its occurrence ^39^. A previous study showed that sialic acid played a role in mediating SVV-GFP infectivity, but the mechanism was unclear ^40^. Our results indicated that SVV entry depended on CMAH, suggesting that the expression of Neu5Gc is conducive to SVV infection. Interestingly, compared with other mammals, human *CMAH* has been evolutionarily mutated and inactivated, which means that humans cannot produce Neu5Gc ^41^, and we speculate that this may also be one of the reasons for the different consequences of SVV infection in humans and pigs.

In conclusion, ANTXR1, heparan sulfate and Neu5Gc were shown to act as host factors of virus entry by our screen, and SVV disease prevention and treatment could potentially benefit from these results. These findings provide significant clues to design new vaccines, develop effective drugs, and evaluate a genetic breeding strategy.

## Materials

### Cells and viruses

IBRS-2, PK-15, MDCK, HEK293-T cells and the derived mutant cell lines were cultured in Dulbecco’s Modified Eagle Medium supplemented with 10% fetal bovine serum (FBS), and 50 μg/mL streptomycin. CRL-2843 and the derived mutant cell lines were cultured in Hyclone RPMI-1640 supplemented with 10% FBS and 50 μg/mL streptomycin. All cell lines were tested and judged free of mycoplasma contamination and maintained at 37 °C (5% CO_2_).

SVV strain isolated in National Foot and Mouth Diseases Reference Laboratory, Lanzhou Veterinary Research Institute, Chinese Academy of Agricultural Sciences was used for screening and validation studies. These SVV strains were propagated in IBRS-2 cells, and viral titers were determined using a doubling dilution assay; the titers were denoted as the 50% cell culture infective dose (TCID_50_)/mL, as determined using the Reed-Muench method. The titrated viruses were preserved at −80 °C (TCID_50_=7.89).

To examine the susceptibilities of cells to SVV, the cells were plated 1 to 2 days prior to inoculation, and then challenged with SVV at the indicated MOI (using the basic medium for dilution) at 37 °C for appropriate time. The viral infection state was evaluated for subsequent analysis.

### Screen library construction

A pooled library encompassing 93,859 different sgRNAs targeting 22,707 pig genes was designed by the Wu laboratory, China Agricultural University, reference genome Sscrofa10.2 from https://www.ensembl.org. The sgRNAs was packaged using a lentivirus library by GenScript, NanJing, China. 1.2×10^8^ IBRS-2 cells were transduced with lentiviruses at a multiplicity of infection (MOI = 0.2). After 2 μg/mL puromycin selection for 3 days, cells were challenged with SVV (MOI = 1) for 24 h. The experiments were performed in three independent experiments. Genomic DNA was extracted from the uninfected cells or the surviving cells. Using Specific primers located on both sides of the sgRNA frame, sgRNA sequences were amplified two rounds, and subjected to next generation sequencing. A lentivirus carried a non-sense sgRNA was transduced into 1×10^7^ as a control cell library. All the used sgRNA amplified primers in this study are listed in **Table S2**.

### Establishment of single gene mutant cell

The three plasmids were transfected into cell (jetPRIME, polyplus) in combination to manufacture a single gene mutant cell line - a plasmid expressing Cas9 protein regulated by 4 μg/mL doxmycin with PB transposon, a plasmid connecting sgRNA targeting gene with PB transposon and puromycin screen marker, and a vector expressing transposase. Genomic DNA was extracted from samples using the DNeasy Blood & Tissue Kit (Qiagen) and positive cells were picked. All the used sgRNA sequence and gene targeting site amplified primers in this study are listed in **Table S3** and **Table S4**.

### Overexpression experiments

The cDNA of *ANTXR1* was amplified from PK-15 cells, The cDNA of *CMAH* was amplified from pig kidney issues and cloned into pcDNA3.1(+)-Myc vector (Invitrogen, Carlsbad, CA, USA) to yield the C-terminal Myc-tagged expression construct. All the used cDNA amplified primers in this study are listed in **Table S5**.

### Western blotting

Cells were lysed using an immunoprecipitation (IP) lysis buffer containing protease inhibitors (Biotechnology, China). The protein concentrations of the extracts were measured with a BCA assay (Beyotime, P0012). Equal amounts of protein from each sample were analyzed by 10% sodium dodecyl sulfate-polyacrylamide gel electrophoresis (SDS-PAGE) and transferred onto polyvinylidene difluoride (PVDF) membranes, which were blocked with 5% skimmed milk and 0.5% Tween-20 in Tris-buffered saline (TBST) at room temperature for one hour and incubated with primary antibodies at 4 °C. The Myc-tagged proteins were probed with a mouse anti-Myc tag monoclonal IgG antibody (1:1000 dilution, CST). The anti-VP2 polyclonal antibody was prepared in our laboratory. Subsequently, the membranes were incubated with a goat anti-rabbit IgG secondary antibody (1:10,000 dilution, Beyotime) or a goat anti-mouse IgG secondary antibody (1:10,000 dilution, Beyotime) for one hour at room temperature, followed by washes in TBST. Proteins were visualized with chemiluminescence and normalized by β-tubulin or β-actin.

### Indirect immunofluorescence assay

To detect SVV infection in mutant cells, an indirect immunofluorescence assay (IFA) was performed. Cells were cultured in 12-well plates and challenged by SVV (MOI = 1). At 8 hpi (hours post infection), the cells were fixed with 4% paraformaldehyde for 30 min and then permeabilized with 0.25% Triton X-100 at room temperature for 5 min. After three times washes with phosphate-buffered saline (PBS), the cells were blocked with 5% bovine serum albumin in PBS for 30 min. Thereafter, the cells were incubated with an anti-VP2 primary antibody overnight at 4 °C. The fluorochrome-conjugated secondary antibody was added in the dark for 6 h at 4 °C. After that, the cells were stained with 4’6’-diamidino-2-phenylindole (DAPI) for 10 min to reveal the nuclei. The fluorescence was visualized using a EVOS M500I imaging System (Invitrogen).

### RNA extraction and quantitative real-time PCR (qRT-PCR)

Total RNA was extracted with TRIzol (Magen) according to the manufacturer’s instructions. G490 5 × All-In-One RT MasterMix (abm) was used for reverse transcription according to the manufacturer’s protocol (Promega). The qRT-PCR analysis was performed in 96-well plates using the BIO-RAD CFX96 detection system. The relative expression level of these genes was calculated using the 2^-ΔΔct^ method, and GAPDH (**glyceraldehyde-3-phosphate dehydrogenase**) mRNA was used as the endogenous control. qRT-PCR was performed on each sample in triplicate. All the used qPCR primers in this study are listed in **Table S6**.

### Coimmunoprecipitations

HEK293-T cells grown in 100 mm plates were transiently co-transfected with 8 µg of Myc-ANTXR1 and 8 µg of Flag-SVV-structure-protein plasmid. The transfectants were harvested 24 h after transfection and subjected to immunoprecipitation assay. The cells were lysed in NP-40 lysis buffer (1% NP-40, 50 mM Tris (pH 8.0), 5 mM EDTA, 150 mM NaCl, 2 mg/mL leupeptin, 2 mg/mL aprotinin, 1 mM phenylmethane sulfonyl fluoride) 30 min, 400 μL lysate incubated with anti-Myc antibody or 400 μL with control IgG antibodies, and 40 μL protein G agarose beads, then placed on a rotating wheel overnight at 4 °C. The agarose beads were pelleted and washed three times in NP-40 lysis buffer. Antibody–antigen complexes bound to the beads were eluted in SDS-PAGE sample buffer by boiling, resolved by SDS-PAGE, and analyzed by Western blotting analysis with the appropriate antibodies.

### Blocking experiments with heparin sodium and an anti-Neu5Gc antibody

SVV was added (MOI = 1) to PK-15 cells after treatment with soluble heparin sodium (Yuanye) at 0.125 mg/mL, 0.25 mg/mL, 0.5 mg/mL, 1 mg/mL, and 2 mg/mL with PK-15 cells for 1 h, respectively.

An anti-Neu5Gc antibody (BioLegend) was preincubated at 1:200 dilution with IBRS-2 cells for 1h, after which SVV was added (MOI = 1). After 1h, media was replaced by DMEM. At 12 hpi, SVV mRNAs was determined using a qPCR assay.

### Statistical analysis

GraphPad Prism software (GraphPad, San Diego, CA) was used to analyze the data. Unpaired two-tailed t test were performed where applicable to determine statistical significance, and the results are presented as the mean ± SEM (\**p* < 0.05, \*\**p* < 0.01, and \*\*\**p* < 0.001).

## Acknowledgements

This work was supported by the National Key Research and Development Program of China (Grant No.2021YFA0805905), the 2020 Research Program of Sanya Yazhou Bay Science and Technology City (202002011), the Plan 111 (B12008), the National Natural Science Foundation of China (32002180), the National Key Research and Development Program of China (Grant No. 2021YFA0805900), Gansu Provincial Major project for science and technology development (21ZD3NA001 and 20ZD7NA006), the Chinese Academy of Agricultural Science and Technology Innovation Project (CAAS-ZDRW202006), and Chinese Universities Scientific Fund.

## Contributions

H.H., X.D., Z.Z., S.W., Y.Z. and H.Z. designed and supervised the project. Y.Z., H.Z., W.T. and W.Y. wrote the paper, with contributions from all authors. W.T. and X.Q. designed the CRISPR/Cas9 library. W.T. performed the screen and verification of gene function. Y.C. assisted with the establishment of single gene mutant cell and data analyses. F.G. performed coimmunoprecipitations. W.T. and K.L. performed indirect immunofluorescence assay.

## Supplementary materials

**Supplement figure 1.**
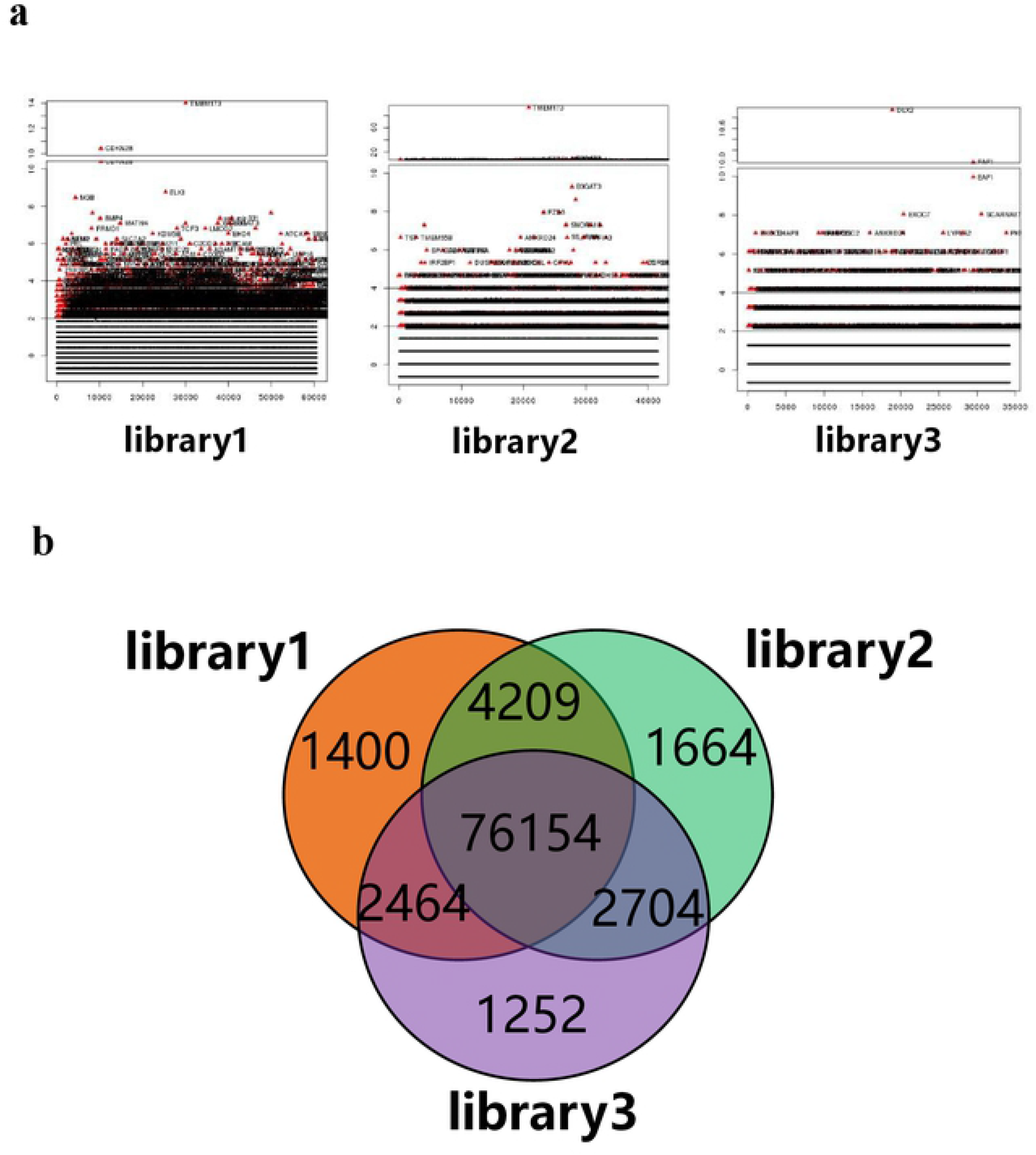
Detailed library sequencing information. **a Enriched genes obtained from three parallel screen libraries.** The target genes were represented by red dots. The abscissa represented genes and the ordinate represented the Z score corresponding to the gene. **b Venn diagrams showed the repeatability of sgRNA enrichment in three parallel screen libraries.** The circles represented the sgRNAs number obtained by deep sequencing.

**Supplement figure 2.**
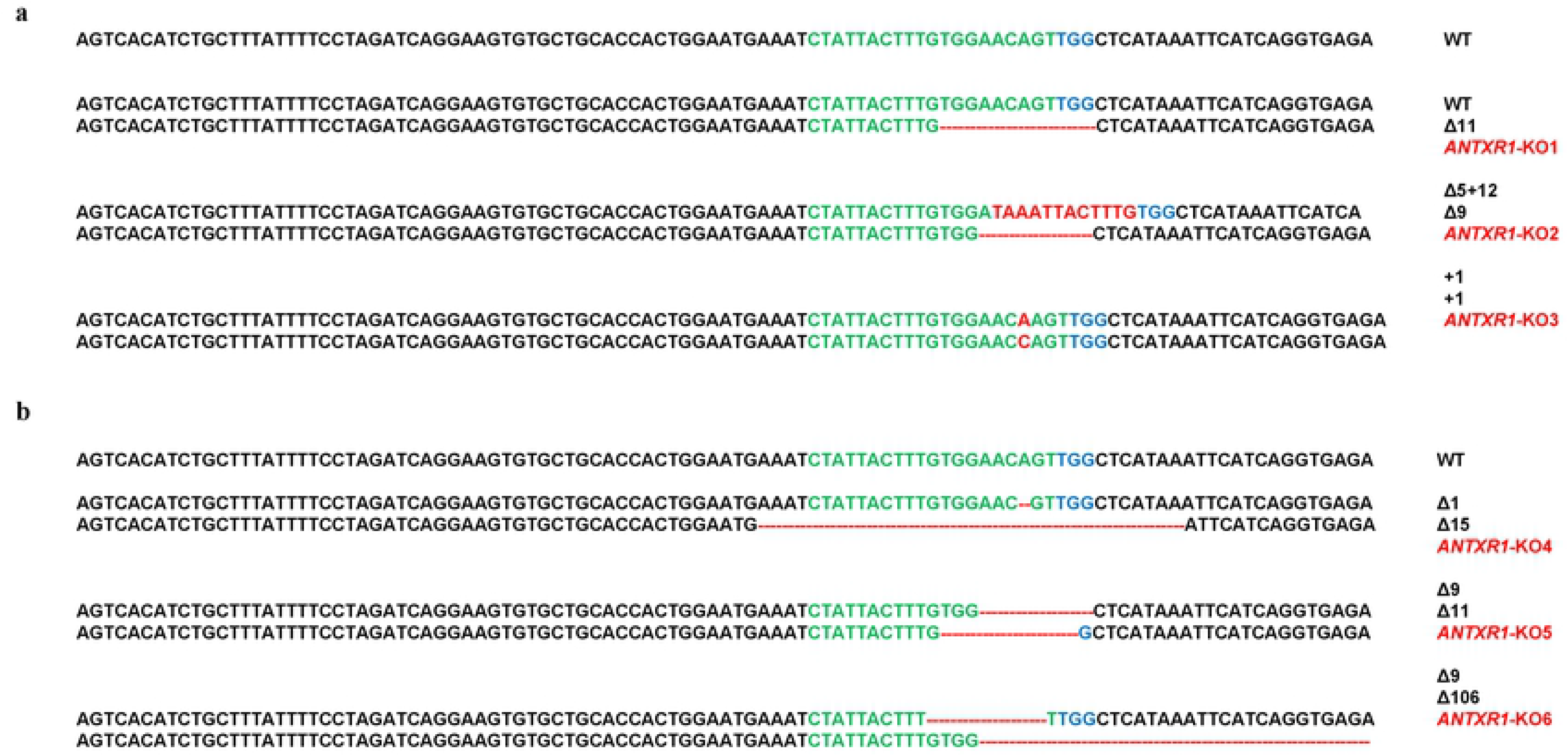
The genomic sequence alternations in *ANTXR1*-KO monoclonal cells. **a The genomic sequence alternations in *ANTXR1*-KO PK15 monoclonal cells.** *ANTXR1*-KO1, *ANTXR1*-KO2 and *ANTXR1*-KO3 monoclonal PK-15 cells were generated by the CRISPR/Cas9 based approach. The sgRNA targeting sites were indicated in green letters and PAM site was indicated in blue letters. The alternations of bases highlighted in red. **b The genomic sequence alternations in *ANTXR1*-KO CRL-2843 monoclonal cells.** *ANTXR1*-KO4, *ANTXR1*-KO5 monoclonal cells, and *ANTXR1*-KO6 CRL-2843 monoclonal cells generated by the CRISPR/Cas9 based approach. The sgRNA targeting sites were indicated in green letters and PAM site was indicated in blue letters. The alternations of bases highlighted in red.

**Supplement figure 3.**
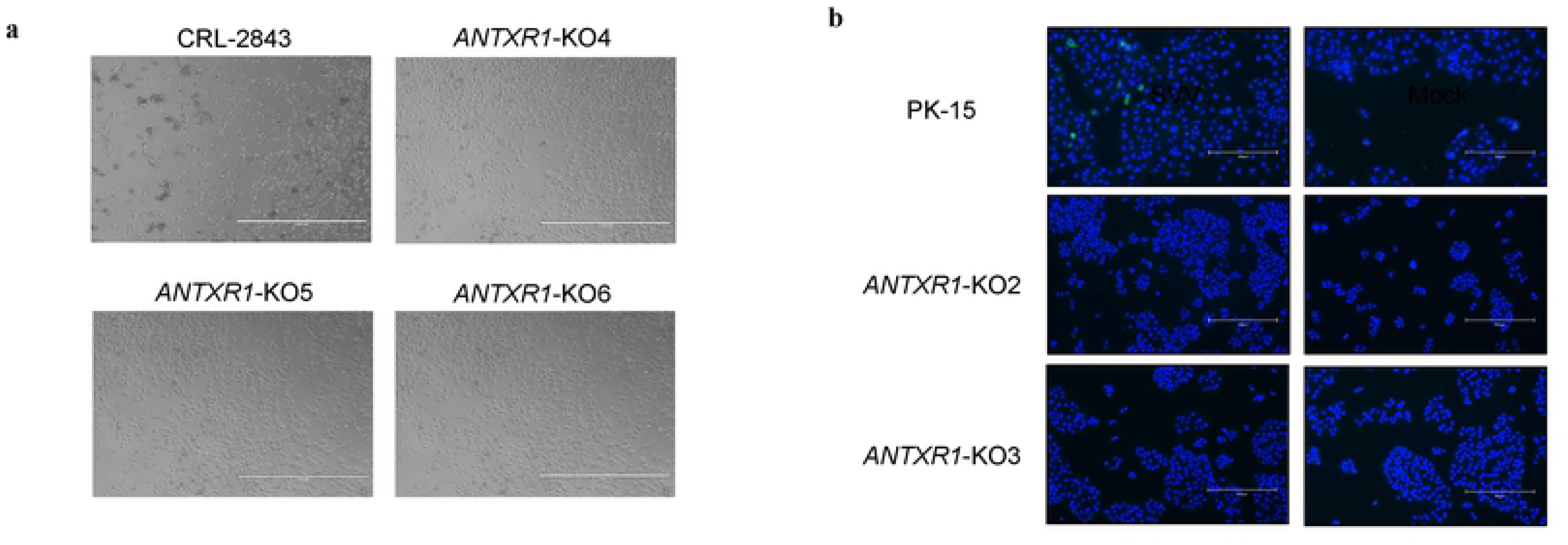
Knockout of ANTXR1 reduced SVV infection. **a CPE of *ANTXR1-*knockout and wild type cells visualized by microscope.** Wild type CRL-2843 cells and *ANTXR1* KO cells were cultured in 12 well plates, and infected with SVV (MOI = 1). At 12 hpi, significant cytopathic effect (CPE) was observed in wild type CRL-2843 rather than in *ANTXR1* KO cells. Scale bars: 1,000 μm. **b Indirect immunofluorescence assay.** Wild type PK-15 cells and *ANTXR1* KO cells were cultured in 24 well plates, and infected with SVV (MOI = 1). At 8 hpi, VP2 protein expression (green) was detected by indirect immunofluorescence assay. Cell nuclei were stained with a NucBlue Live Ready Probe (blue). Scale bars: 300 μm.

**Supplement figure 4.**
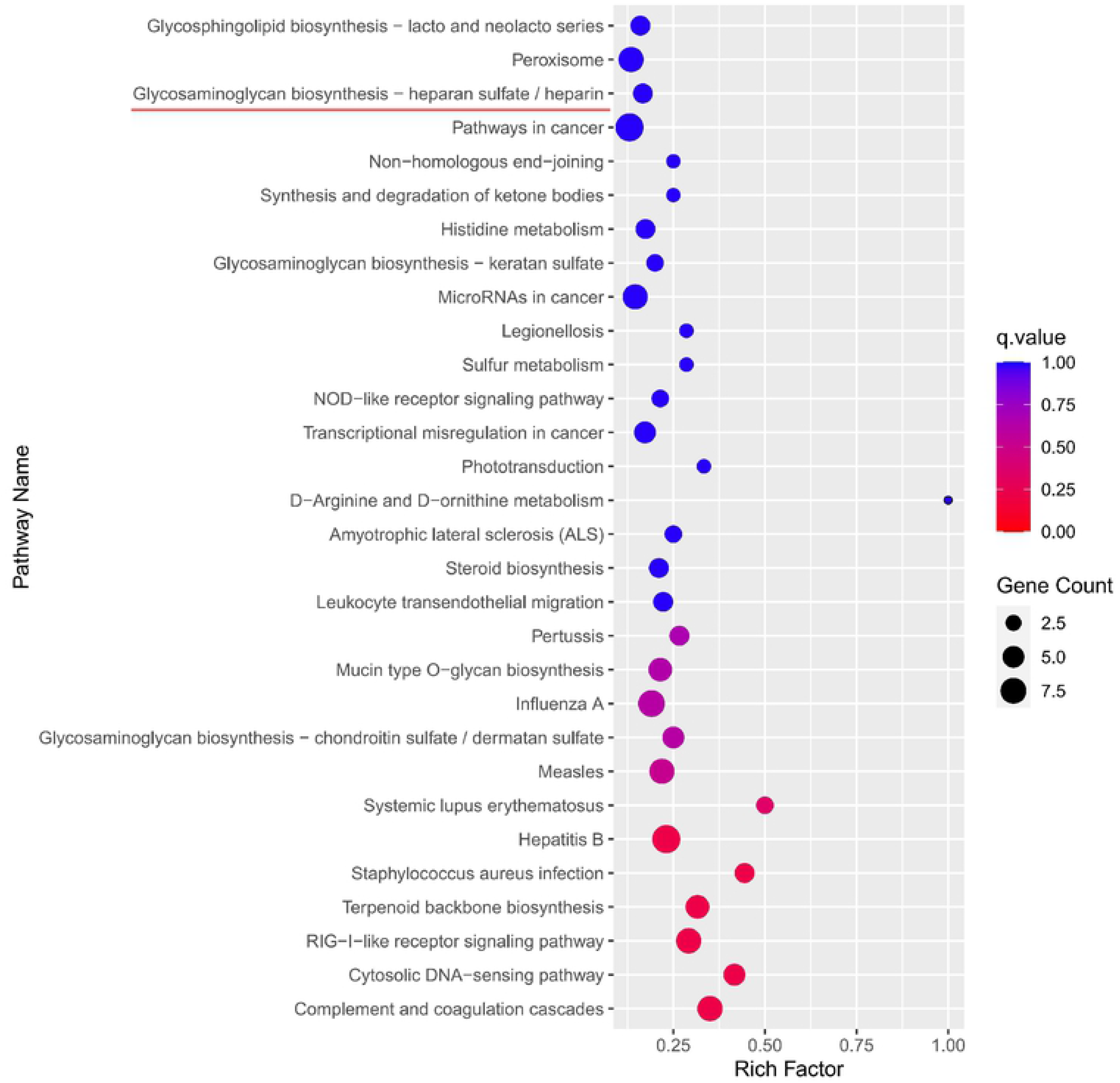
KEGG analysis of RNA-seq results. Wild type PK-15 cells were cultured in six well plates and infected with SVV (MOI = 1). At 12 hpi, the RNA of cells were collected for RNA-seq. Compared to non-infected PK-15 cells, top 30 enrichment pathways in PK-15 cells infected with SVV (MOI = 1) were displayed.

**Supplement figure 5.**
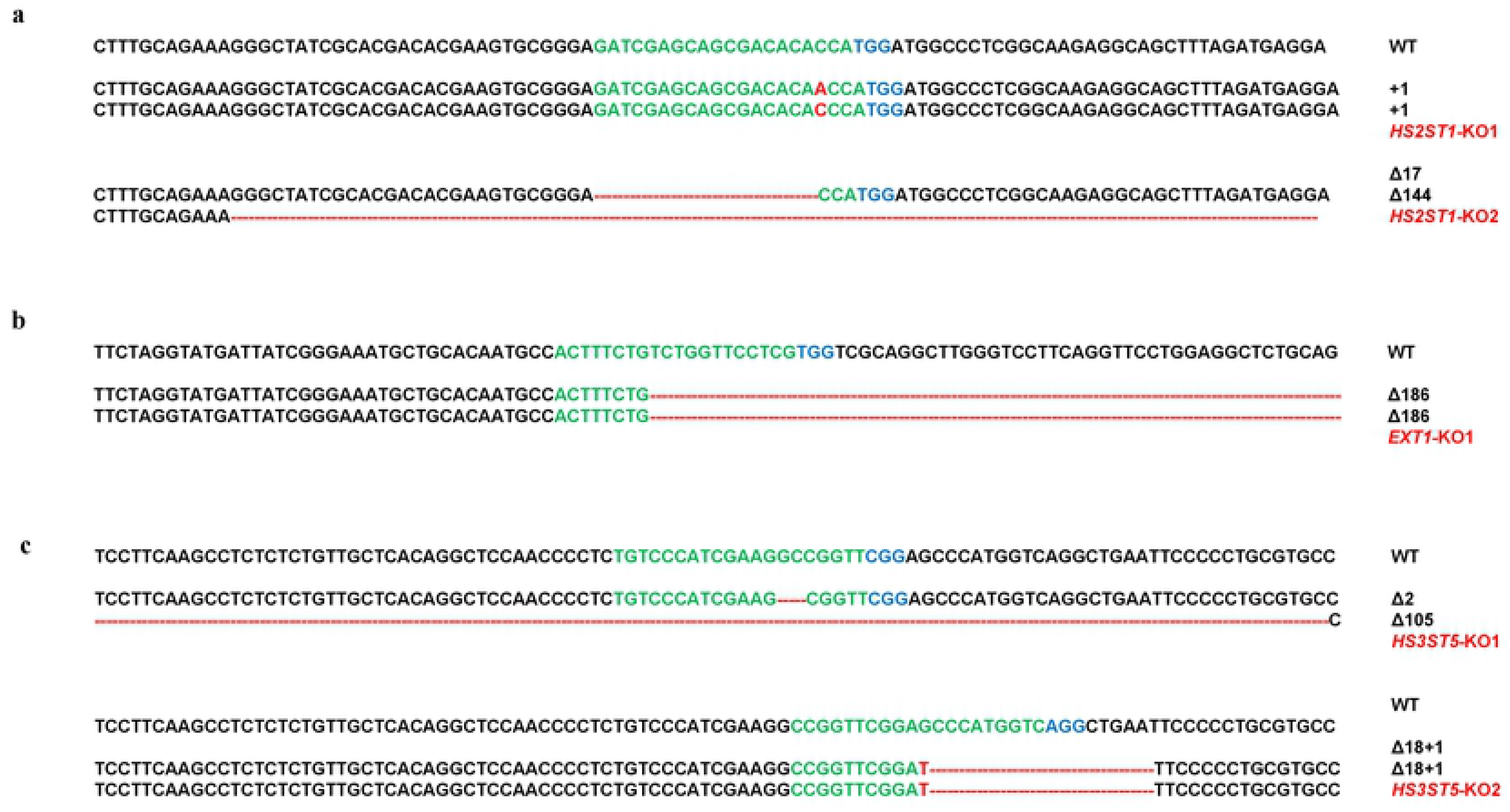
The genomic sequence alternations in *HS2ST1*, *EXT1*, *HS3ST5* knockout monoclonal cells. **a The genomic sequence alternations in *HS2ST1*-KO PK15 monoclonal cells.** Both monoclonal cells were generated by the CRISPR/Cas9 based approach. The sgRNA targeting sites were indicated in green letters and PAM site was indicated in blue letters. The alternations of bases highlighted in red. **b The genomic sequence alternations in EXT1-KO PK15 monoclonal cell.** Monoclonal cell was generated by the CRISPR/Cas9 based approach. The sgRNA targeting sites were indicated in green letters and PAM site was indicated in blue letters. The alternations of bases highlighted in red. **c The genomic sequence alternations in HS3ST5-KO PK15 monoclonal cells.** Both monoclonal cells were generated by the CRISPR/Cas9 based approach. The sgRNA targeting sites were indicated in green letters and PAM site was indicated in blue letters. The alternations of bases highlighted in red.

**Supplement figure 6.**
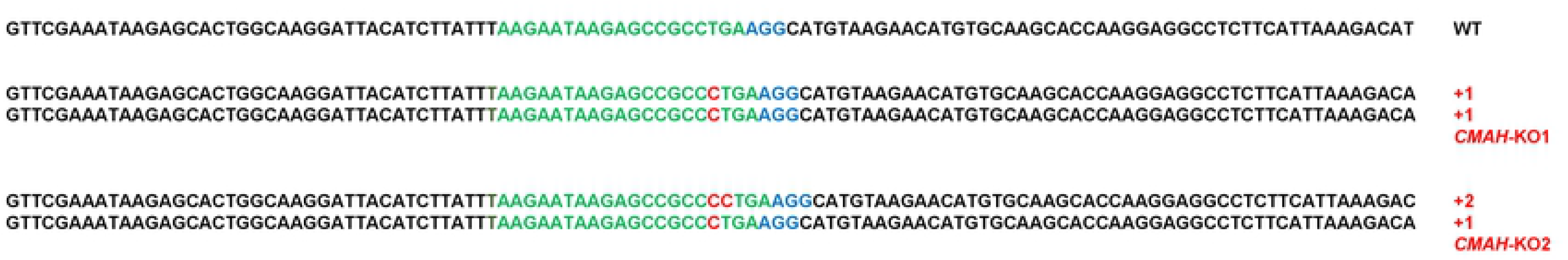
The genomic sequence alternations in *CMAH*-KO monoclonal cells. Both monoclonal cells were generated by the CRISPR/Cas9 based approach. The sgRNA targeting sites were indicated in green letters and PAM site was indicated in blue letters. The alternations of bases highlighted in red.

**Table S1.**
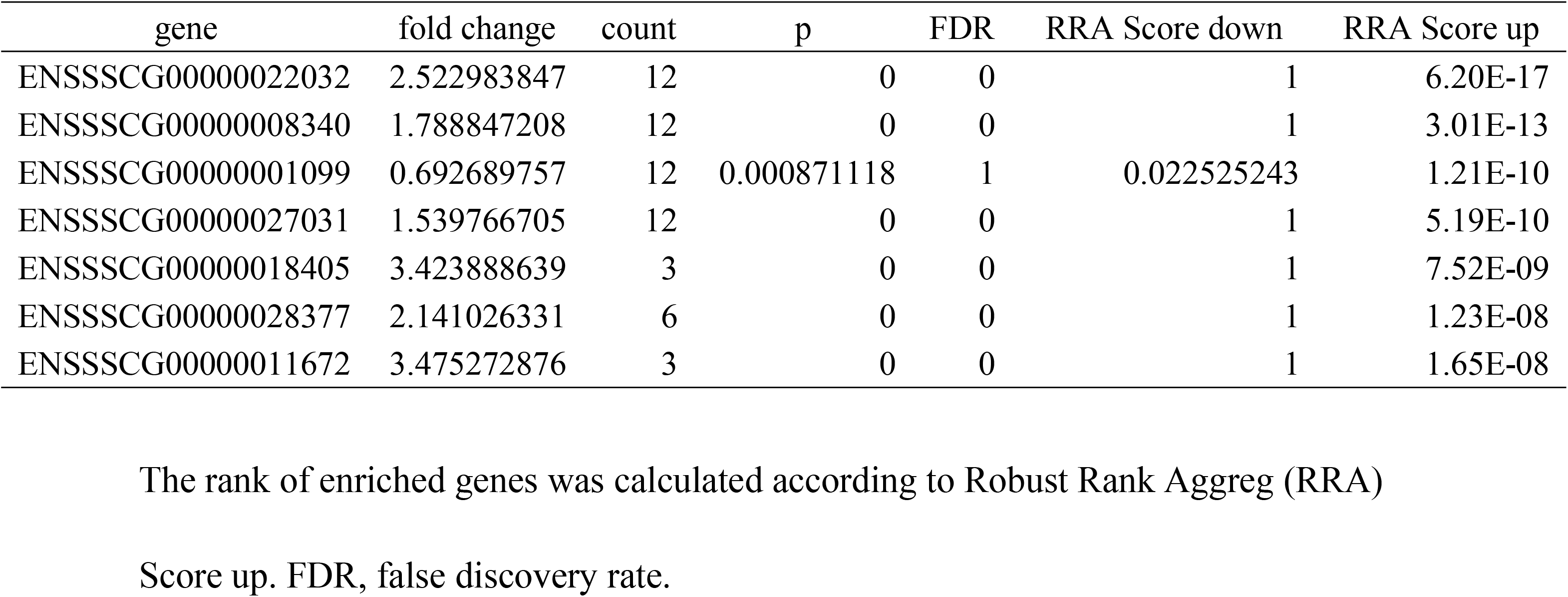
The top 7 candidate genes of screen.

**Table S2.**
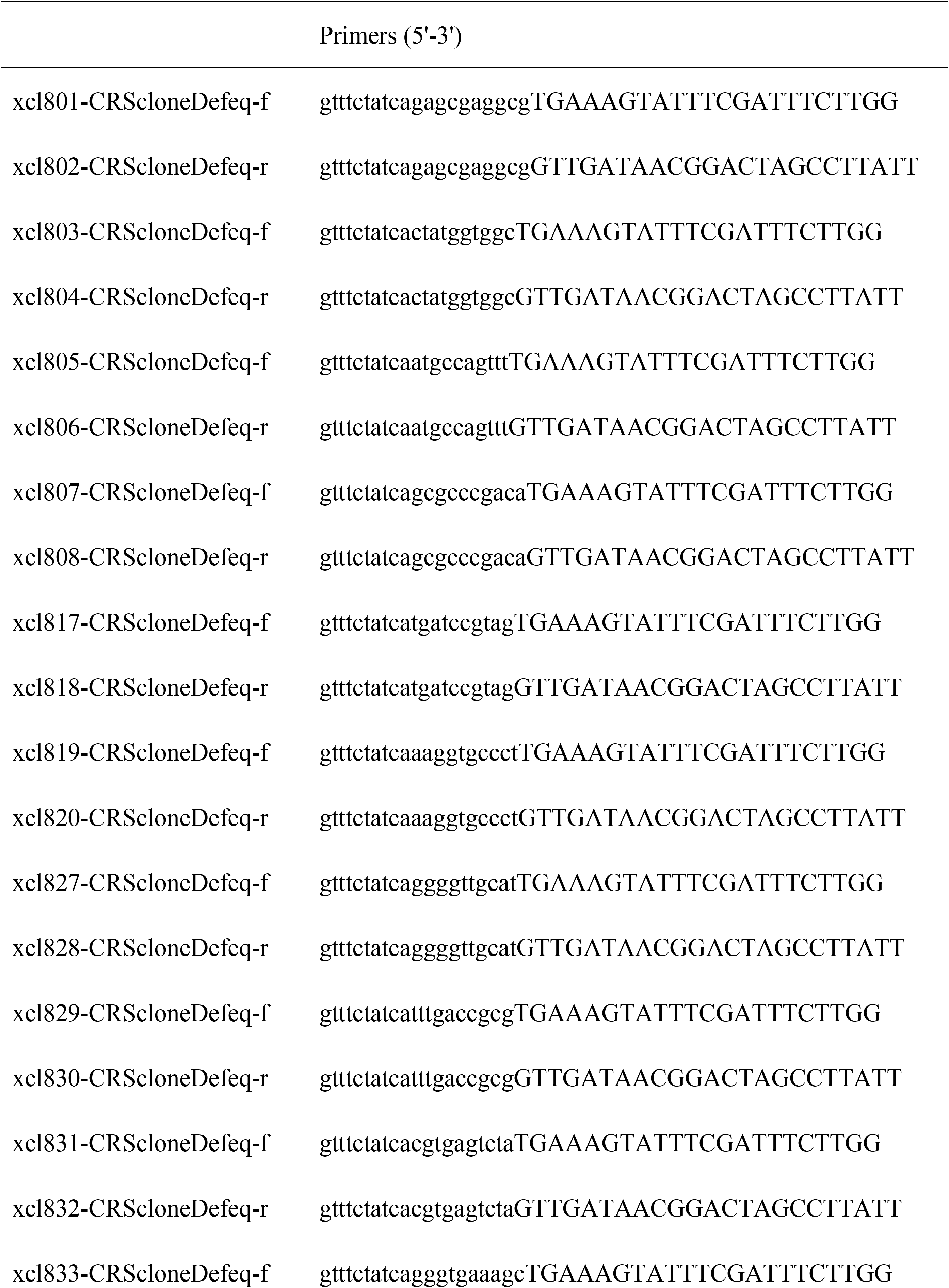
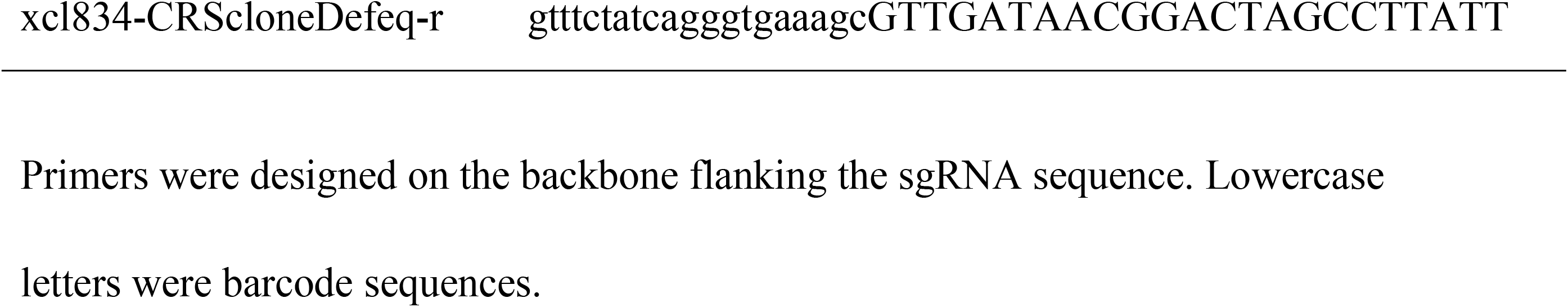
The sgRNA amplified primers used in this study.

**Table S3.**
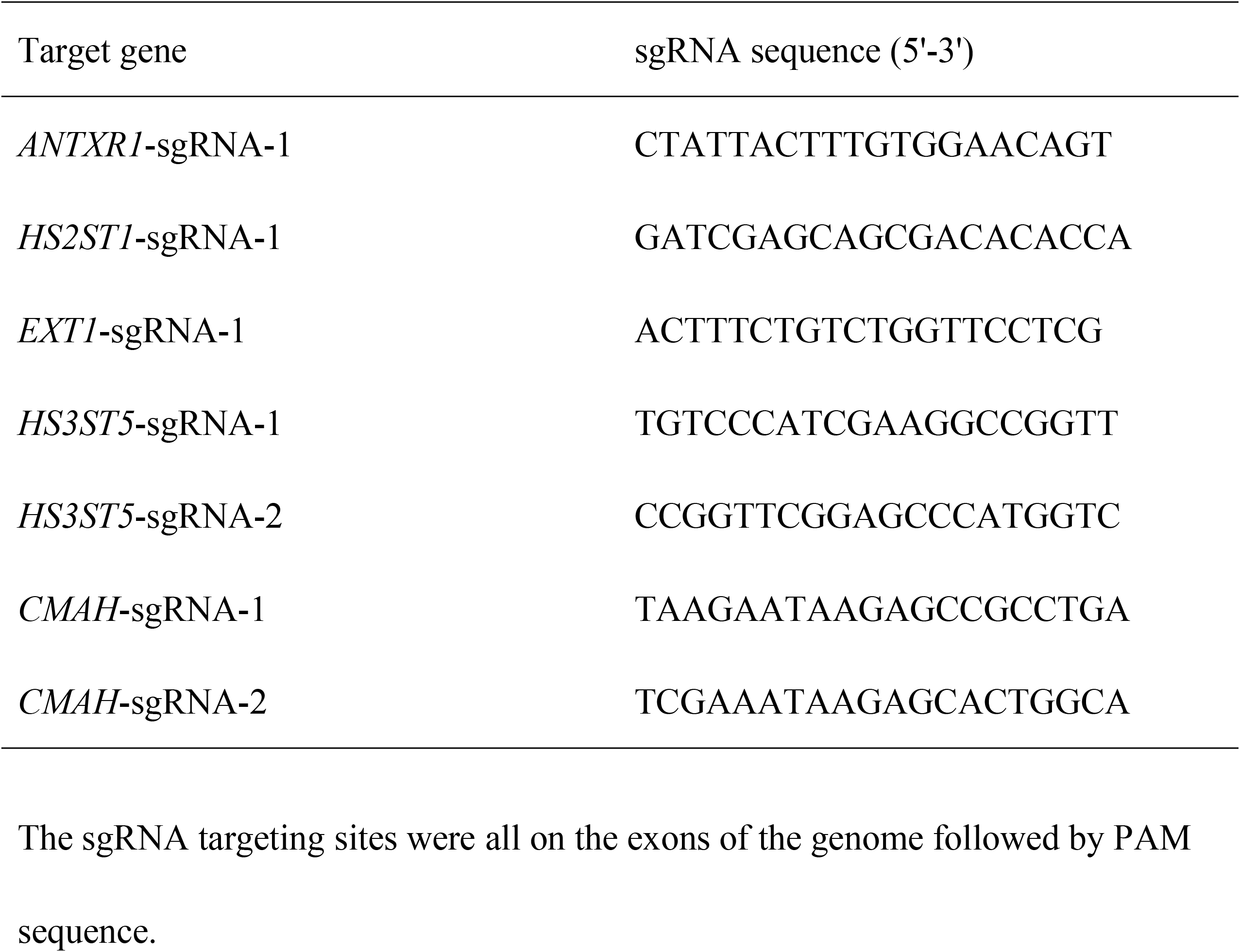
The sgRNA sequence used in this study.

**Table S4.**
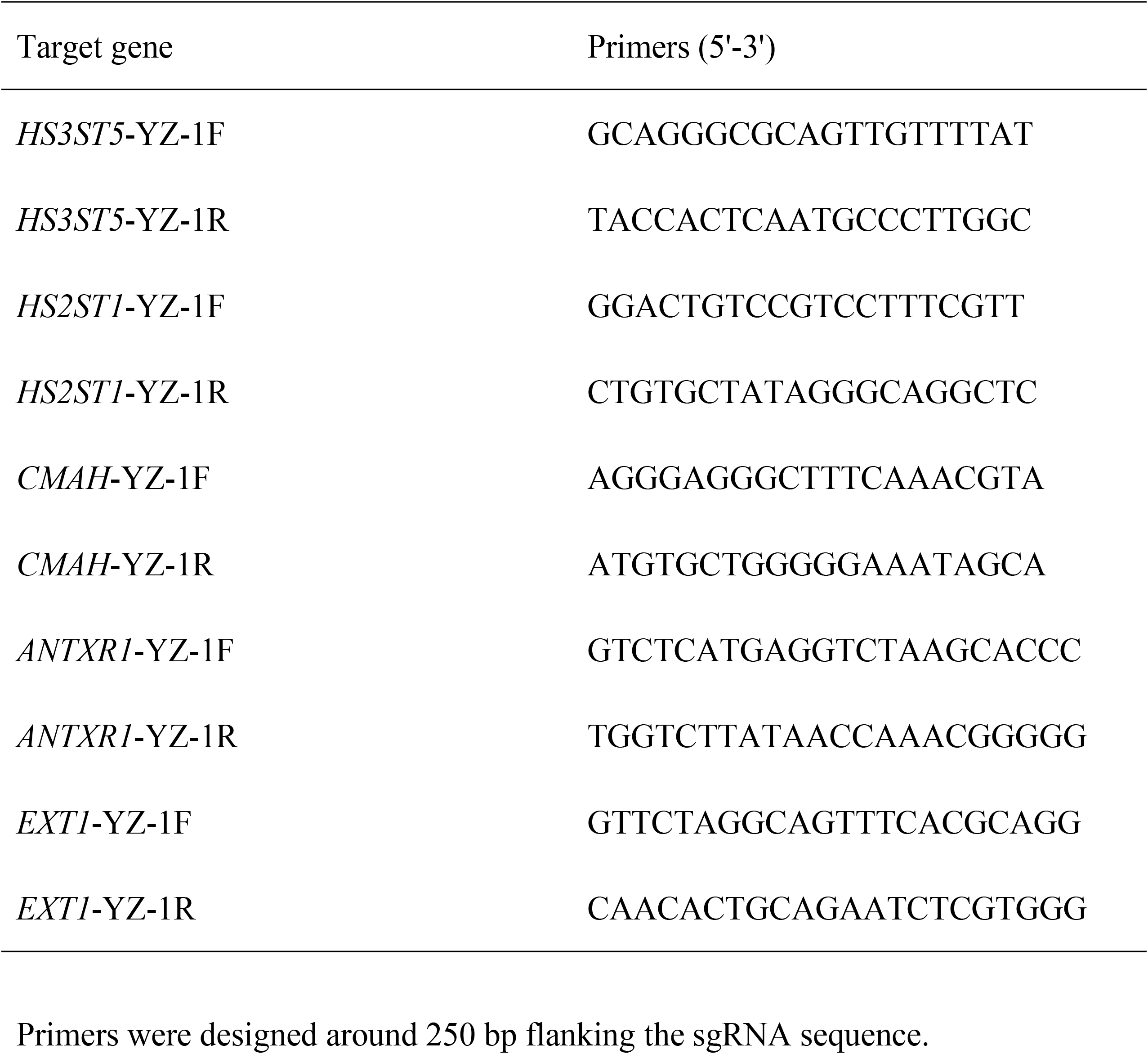
The gene targeting site amplified primers used in this study.

**Table S5.**
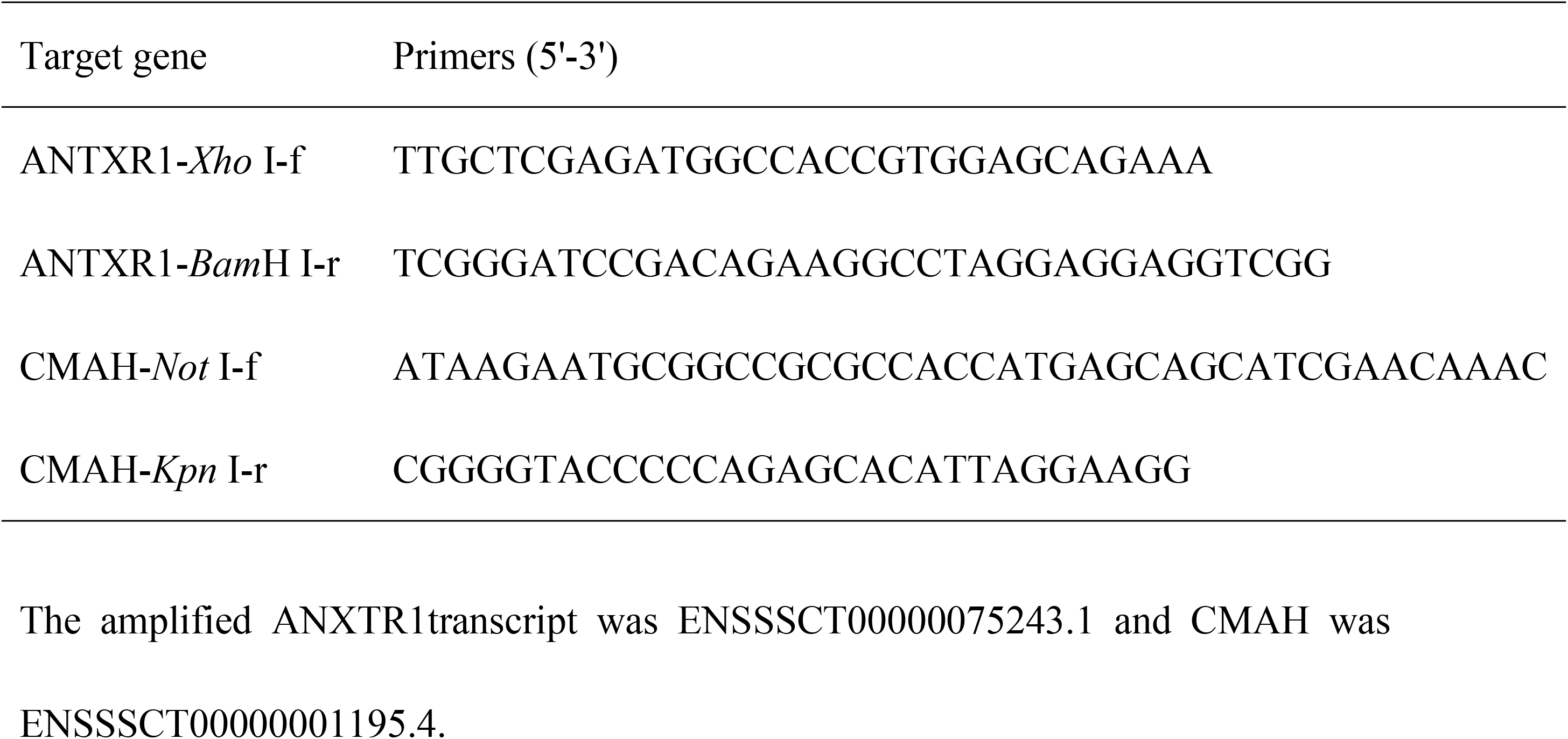
The cDNA amplified primers used in this study.

**Table S6.**
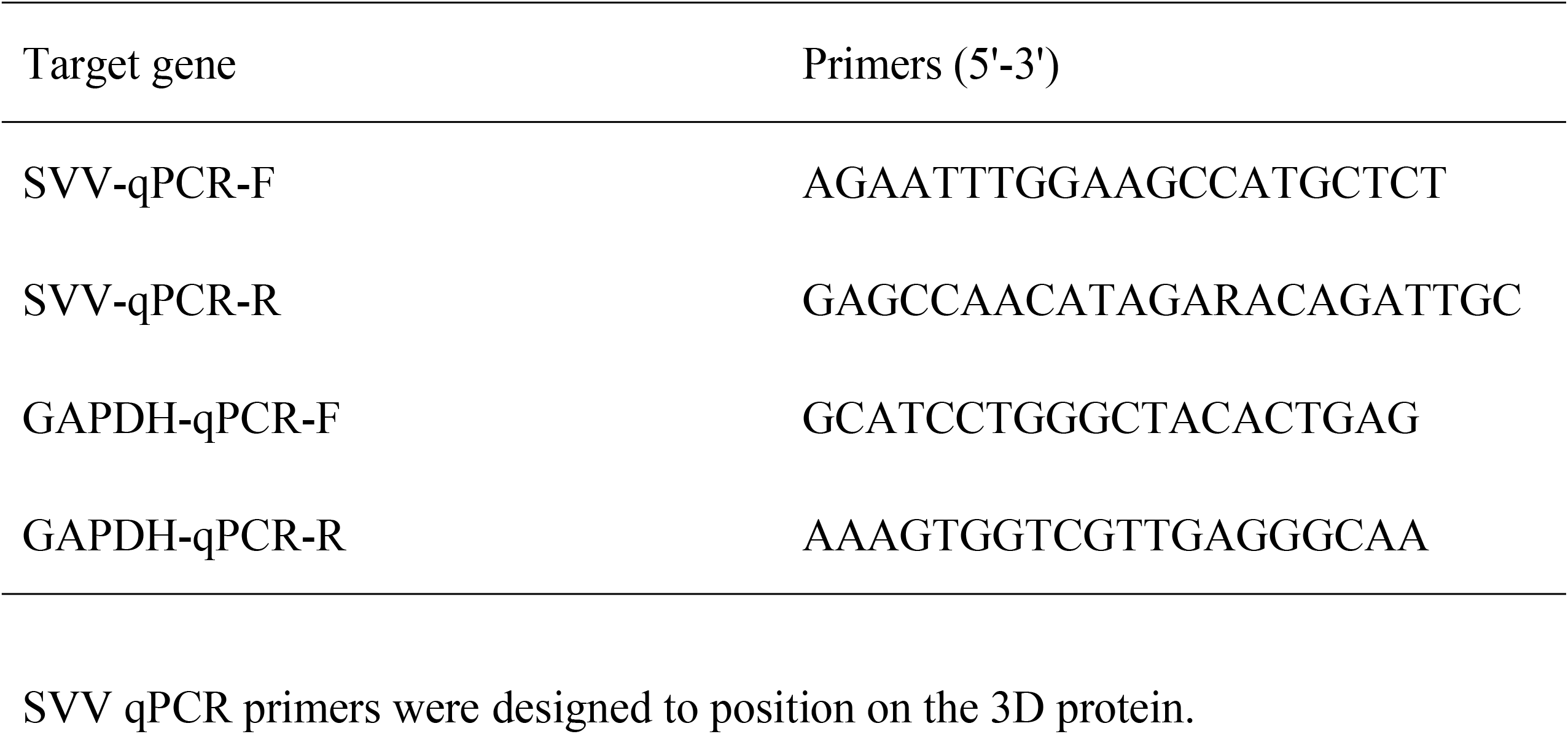
The qPCR primers used in this study.

## Reference

1. Canning, P. et al. Neonatal mortality, vesicular lesions and lameness associated with Senecavirus A in a U.S. Sow Farm. Transbound Emerg. Dis. 63, 373–378 (2016).

2. Guo, B. et al. Novel Senecavirus A in swine with vesicular disease, United States, July 2015. Emerg. Infect. Dis. 22, 1325–1327 (2016).

3. Hause, B. M., Myers, O., Duff, J. & Hesse, R. A. Senecavirus A in pigs, United States, 2015. Emerg. Infect. Dis. 22, 1323–1325 (2016).

4. Houston, E. et al. Seroprevalence of Senecavirus A in sows and grower-finisher pigs in major swine producing-states in the United States. Prev. Vet. Med. 165, 1–7 (2019).

5. Leme, R. A. et al. Clinical manifestations of Senecavirus A infection in neonatal pigs, Brazil, 2015. Emerg. Infect. Dis. 22, 1238–1241 (2016).

6. Leme, R. A., Oliveira, T. E. S., Alfieri, A. F., Headley, S. A. & Alfieri, A. A. Pathological, immunohistochemical and molecular findings associated with Senecavirus A-induced lesions in neonatal piglets. J. Comp. Pathol. 155 (2016).

7. Leme, R. A. et al. Senecavirus A: An emerging vesicular infection in Brazilian pig herds. Transbound. Emerg. Dis. 62, 603–611 (2015).

8. Oliveira, T. E. S. et al. Histopathological, immunohistochemical, and ultrastructural evidence of spontaneous Senecavirus A-induced lesions at the choroid plexus of newborn piglets. Sci. Rep. 7, 16555 (2017).

9. Qian, S., Fan, W., Qian, P., Chen, H. & Li, X. Isolation and full-genome sequencing of Seneca Valley virus in piglets from China, 2016. Virol. J. 13, 173 (2016).

10. Wu, Q. et al. The first identification and complete genome of Senecavirus A affecting pig with idiopathic vesicular disease in China. Transbound Emerg. Dis. 64, 1633–1640 (2017).

11. Li, W. et al. Angiotensin-converting enzyme 2 is a functional receptor for the SARS coronavirus. Nature 426, 450–454 (2003).

12. Chan, K. K. et al. Engineering human ACE2 to optimize binding to the spike protein of SARS coronavirus 2. Science 369, 1261–1265 (2020).

13. Proudfoot, N., Mcfarlane, N., Whitelaw, N. & Lillico, N. Livestock breeding for the 21st century: the promise of the editing revolution. Front. Agr. Sci. Eng. 7, 7 (2020).

14. Whitworth, K. M. et al. Gene-edited pigs are protected from porcine reproductive and respiratory syndrome virus. Nat. Biotechnol. 34, 20–22 (2016).

15. Burkard, C. et al. Precision engineering for PRRSV resistance in pigs: Macrophages from genome edited pigs lacking CD163 SRCR5 domain are fully resistant to both PRRSV genotypes while maintaining biological function. PLoS. Pathog. 13, e1006206 (2017).

16. Chen, J. et al. Generation of pigs resistant to highly Pathogenic-Porcine Reproductive and Respiratory Syndrome virus through gene editing of CD163. Int. J. Biol. Sci. 15, 481–492 (2019).

17. Hales, L. M. et al. Complete genome sequence analysis of Seneca Valley virus-001, a novel oncolytic picornavirus. J. Gen. Virol. 89, 1265–1275 (2008).

18. Miles, L. A. et al. Anthrax toxin receptor 1 is the cellular receptor for Seneca Valley virus. J. Clin. Invest. 127, 2957–2967 (2017).

19. Wei, J. et al. Genome-wide CRISPR screens reveal host factors critical for SARS-CoV-2 infection. Cell 184, 76–91 e13 (2021).

20. Ganaie, S. S. et al. Lrp1 is a host entry factor for Rift Valley fever virus. Cell 184, 5163–5178 e5124 (2021).

21. Clark, L. E. et al. VLDLR and ApoER2 are receptors for multiple alphaviruses. Nature 602, 475–480 (2022).

22. Santelli, E., Bankston, L. A., Leppla, S. H. & Liddington, R. C. Crystal structure of a complex between anthrax toxin and its host cell receptor. Nature 430, 905–908 (2004).

23. Venkataraman, S. et al. Structure of Seneca Valley Virus-001: an oncolytic picornavirus representing a new genus. Structure 16, 1555–1561 (2008).

24. Broszeit, F. et al. N-glycolylneuraminic acid as a receptor for Influenza A viruses. Cell Rep. 27, 3284–3294 e3286 (2019).

25. Jayawardena, N. et al. Structural basis for anthrax toxin receptor 1 recognition by Seneca Valley Virus. Proc. Natl. Acad. Sci. U. S. A. 115, e10934–e10940 (2018).

26. Cao, L. et al. Seneca Valley virus attachment and uncoating mediated by its receptor anthrax toxin receptor 1. Proc. Natl. Acad. Sci. U. S. A. 115, 13087–13092 (2018).

27. Cullen, M. et al. Host-derived tumor endothelial marker 8 promotes the growth of melanoma. Cancer Res. 69, 6021–6026 (2009).

28. Stranecky, V. et al. Mutations in ANTXR1 cause GAPO syndrome. Am. J. Hum. Genet. 92, 792–799 (2013).

29. Chen, P. R. et al. Disruption of anthrax toxin receptor 1 in pigs leads to a rare disease phenotype and protection from Senecavirus A infection. Sci. Rep. 12, 5009 (2022).

30. Song, R., Wang, Y. & Zhao, J. Base editing in pigs for precision breeding. Front. Agr. Sci. Eng. 7, 161–170 (2020).

31. Kreuger, J. & Kjellen, L. Heparan sulfate biosynthesis: regulation and variability. J. Histochem. Cytochem. 60, 898–907 (2012).

32. Xian, X., Gopal, S. & Couchman, J. R. Syndecans as receptors and organizers of the extracellular matrix. Cell Tissue Res. 339, 31–46 (2010).

33. Lindahl, U. & Li, J. P. Interactions between heparan sulfate and proteins-design and functional implications. Int. Rev. Cell Mol. Biol. 276, 105–159 (2009).

34. Hacker, U., Nybakken, K. & Perrimon, N. Heparan sulphate proteoglycans: the sweet side of development. Nat. Rev. Mol. Cell Biol. 6, 530–541 (2005).

35. Clausen, T. M. et al. SARS-CoV-2 Infection depends on cellular heparan sulfate and ACE2. Cell 183, 1043–1057 e1015 (2020).

36. Volland, A. et al. Heparan sulfate proteoglycans serve as alternative receptors for low affinity LCMV variants. PLoS. Pathog. 17, e1009996 (2021).

37. Tanaka, A. et al. Genome-wide screening uncovers the significance of N-sulfation of heparan sulfate as a host cell factor for Chikungunya virus Infection. J. Virol. 91, e00432–17 (2017).

38. Spruit, C. M. et al. N-glycolylneuraminic acid in animal models for human Influenza A virus. Viruses 13, 815 (2021).

39. Tu, C. F. et al. Lessening of porcine epidemic diarrhoea virus susceptibility in piglets after editing of the CMP-N-glycolylneuraminic acid hydroxylase gene with CRISPR/Cas9 to nullify N-glycolylneuraminic acid expression. PLoS. One 14, e0217236 (2019).

40. Liu, Z. et al. Intravenous injection of oncolytic picornavirus SVV-001 prolongs animal survival in a panel of primary tumor-based orthotopic xenograft mouse models of pediatric glioma. Neuro. Oncol. 15, 1173–1185 (2013).

41. Varki, A. Multiple changes in sialic acid biology during human evolution. Glycoconj. J 26, 231–245 (2009).

